# Between semelparity and iteroparity: empirical evidence for a continuum of modes of parity

**DOI:** 10.1101/107268

**Authors:** P. William Hughes

## Abstract

The number of times an organism reproduces (i.e. its mode of parity) is a fundamental life-history character, and evolutionary and ecological models that compare the relative fitness of strategies are common in life history theory and theoretical biology. Despite the success of mathematical models designed to compare intrinsic rates of increase between annual-semelparous and perennial-iteroparous reproductive schedules, there is widespread evidence that variation in reproductive allocation among semelparous and iteroparous organisms alike is continuous. This paper reviews the ecological and molecular evidence for the continuity and plasticity of modes of parity––that is, the idea that annual-semelparous and perennial-iteroparous life histories are better understood as endpoints along a continuum of possible strategies. I conclude that parity should be understood as a continuum of different modes of parity, which differ by the degree to which they disperse or concentrate reproductive effort in time. I further argue that there are three main implications of this conclusion: (1) That seasonality should not be conflated with parity; (2) that mathematical models purporting to explain the evolution of semelparous life histories from iteroparous ones (or vice versa) should not assume that organisms can only display either an annual-semelparous life history or a perennial-iteroparous one; and (3) that evolutionary ecologists should examine the physiological or molecular basis of traits underlying different modes of parity, in order to obtain a general understanding of how different life history strategies can evolve from one another.

## I. INTRODUCTION

Semelparity (and the related botanical term “monocarpy”) describes the life history defined by a single, highly fecund bout of reproduction, and can be contrasted with iteroparity (“polycarpy”), the life history defined by repeated (i.e. “iterative”) bouts of reproduction throughout life. Identifying the reasons why organisms adopt either mode of parity is one of life history theory’s oldest problems, having been considered by both Aristotle (History of Animals, BkIX, 622 1-30, trans. Thompson, 1907) and Linnaeus (Linnaeus, 1744). In contemporary evolutionary ecology, this problem has been formalized by age-structured demographic models that seek to explain the eco-evolutionary dynamics of reproductive patterns by comparing the intrinsic rates of increase of reproductive strategies (Cole, 1954; Bryant, 1971; Charnov and Schaffer, 1973; Young, 1981; Omielan, 1991; Su and Peterman, 2012; Javoiš, 2013; Vaupel et al., 2013; Cushing, 2015). In such models, two modes of parity are considered, classified by whether they express all reproductive effort in a single year (semelparity), or in more than one (iteroparity). I refer to this simplified conception as the “discrete conception of parity”. The main advantage of the discrete conception of parity is its analytical simplicity; given population growth data, intrinsic rates of increase can be easily computed and directly compared. Some intraspecific comparisons between phenotypically similar semelparous and iteroparous congeners conform to the predictions of demographic models based on the discrete conception of parity (e.g. Fritz et al., 1982; Young, 1984, 1990; Iguchi and Tsukamoto, 2001).

However, in this review I will argue that despite the modest successes—both theoretical and empirical—of evolutionary explanations rooted in the discrete conception of parity, sufficient evidence has been accumulated to make it clear that, like many other life-history traits, parity is a continuous variable, and that semelparity and iteroparity are the endpoints of a continuum of possible strategies that define the distribution of reproductive effort through time, rather than simple alternatives describing whether an organism fatally reproduces in a given year or not. On this account, semelparity can be understood as the strategy defined by concentrating reproductive effort in time, and iteroparity as the strategy defined by distributing reproductive effort over longer timescales. I refer to this idea hereafter as the “continuous conception of parity”. It is important to note that the continuous conception of parity should not be conflated with the related terms “annuality” and “perenniality”. These terms specify strategies defined by the “digitization” of reproduction in response to seasonal effects supervening on the process of reproduction, rather than describing how concentrated reproductive effort is in time. This distinction is further discussed later.

The idea that parity itself is continuous and not discrete is not new (see: Hughes and Simons, 2014c; Kirkendall and Stenseth, 1985; Unwin, Kinnison and Quinn, 1999; Roff, 1992), but to date no systematic exposition of the empirical evidence supporting the different conceptions of parity has yet been undertaken. Furthermore, evolutionary explanations of life-history differences between clades with differing modes of parity continue to rely on the discrete conception of parity (e.g. Lopes and Leiner, 2015), and mathematical models based on the formalization of this assumption continue to be produced (e.g. Benton and Grant, 1999, Davydova et al., 2005; Vaupel et al., 2013). However, because of the ubiquity of evolutionary transitions from iteroparity to semelparity (Table 1), understanding parity as a continuous trait is important for understanding the underlying eco-evolutionary dynamics that affect the fitness of life-history strategies.

**Table 1.**
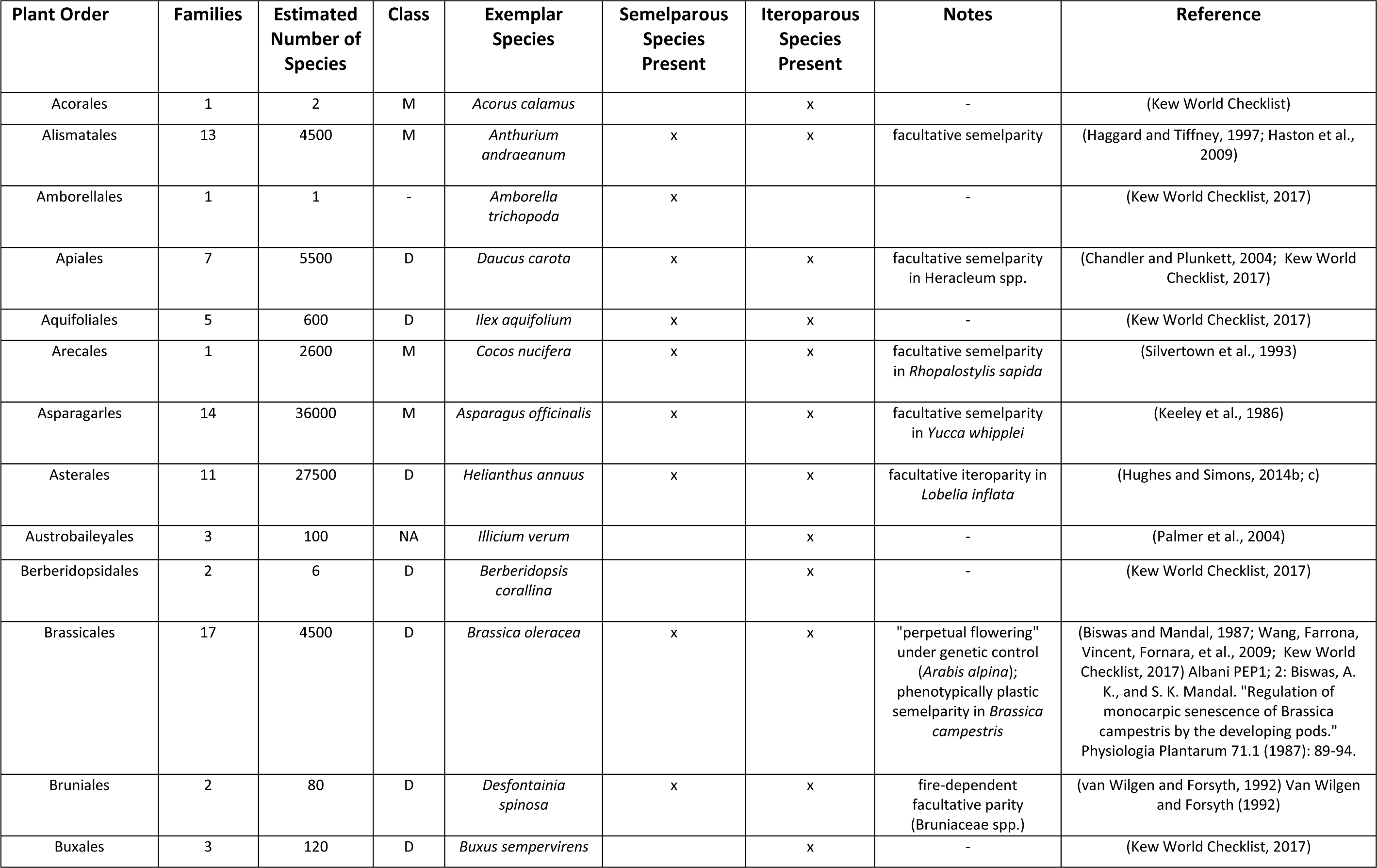
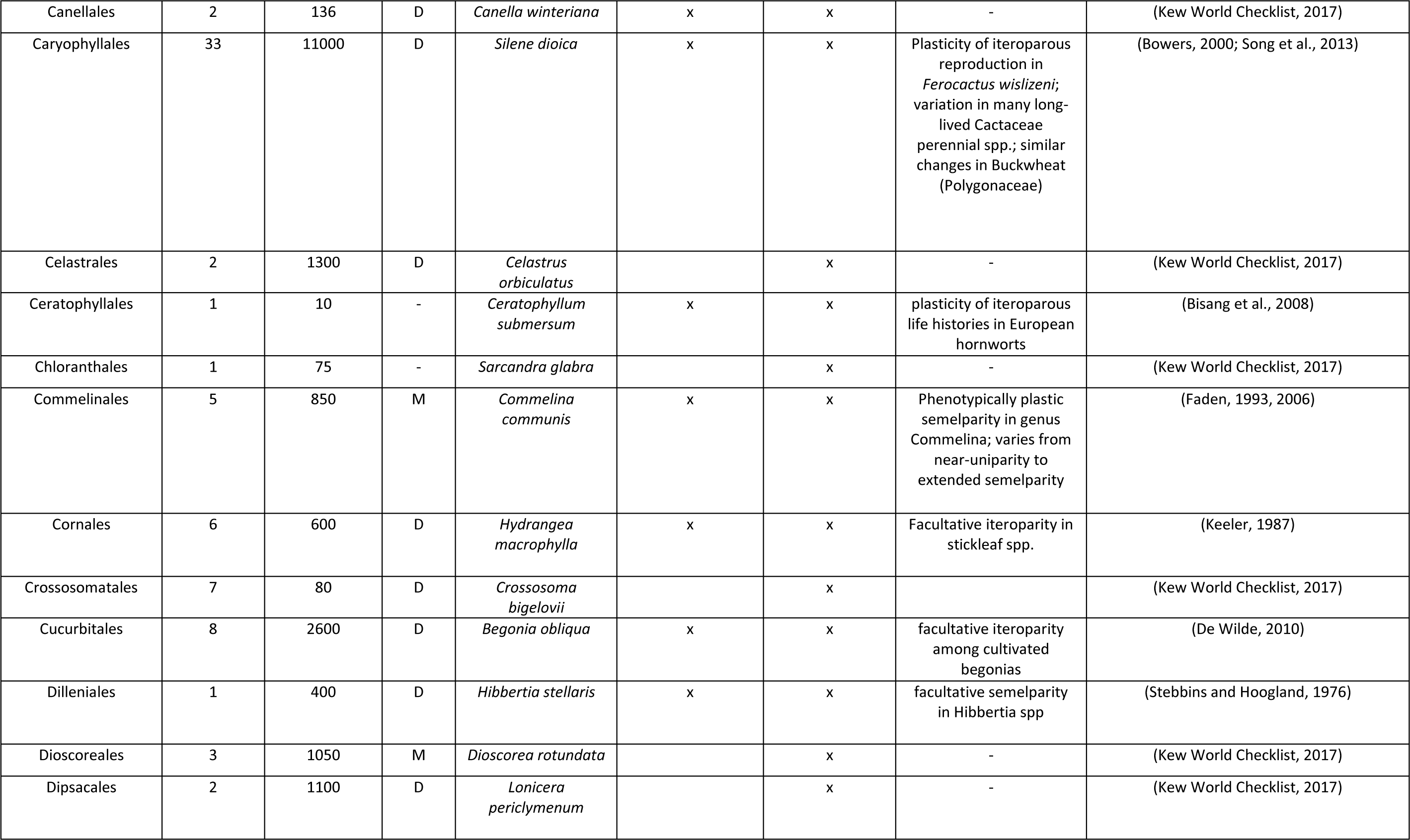
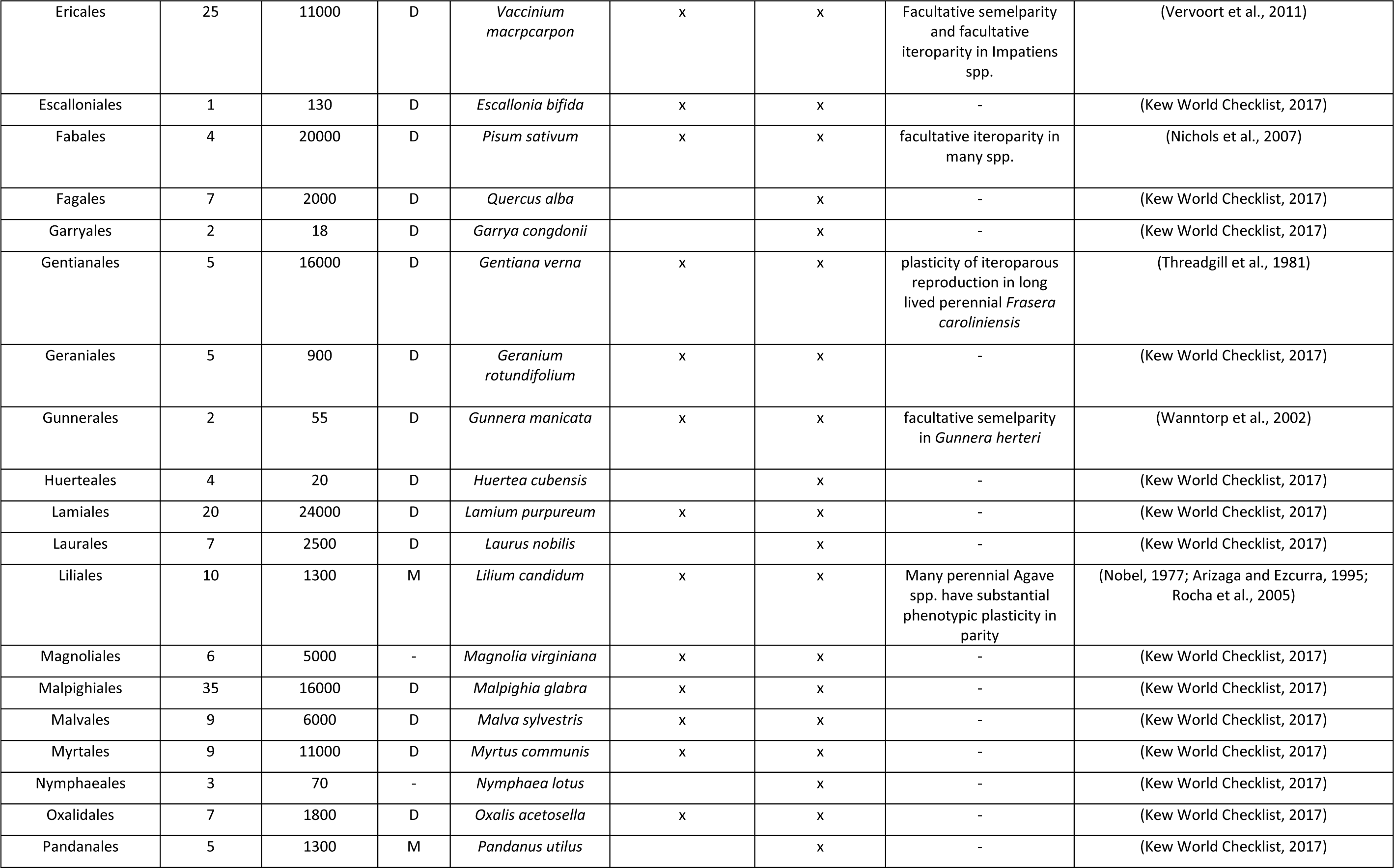
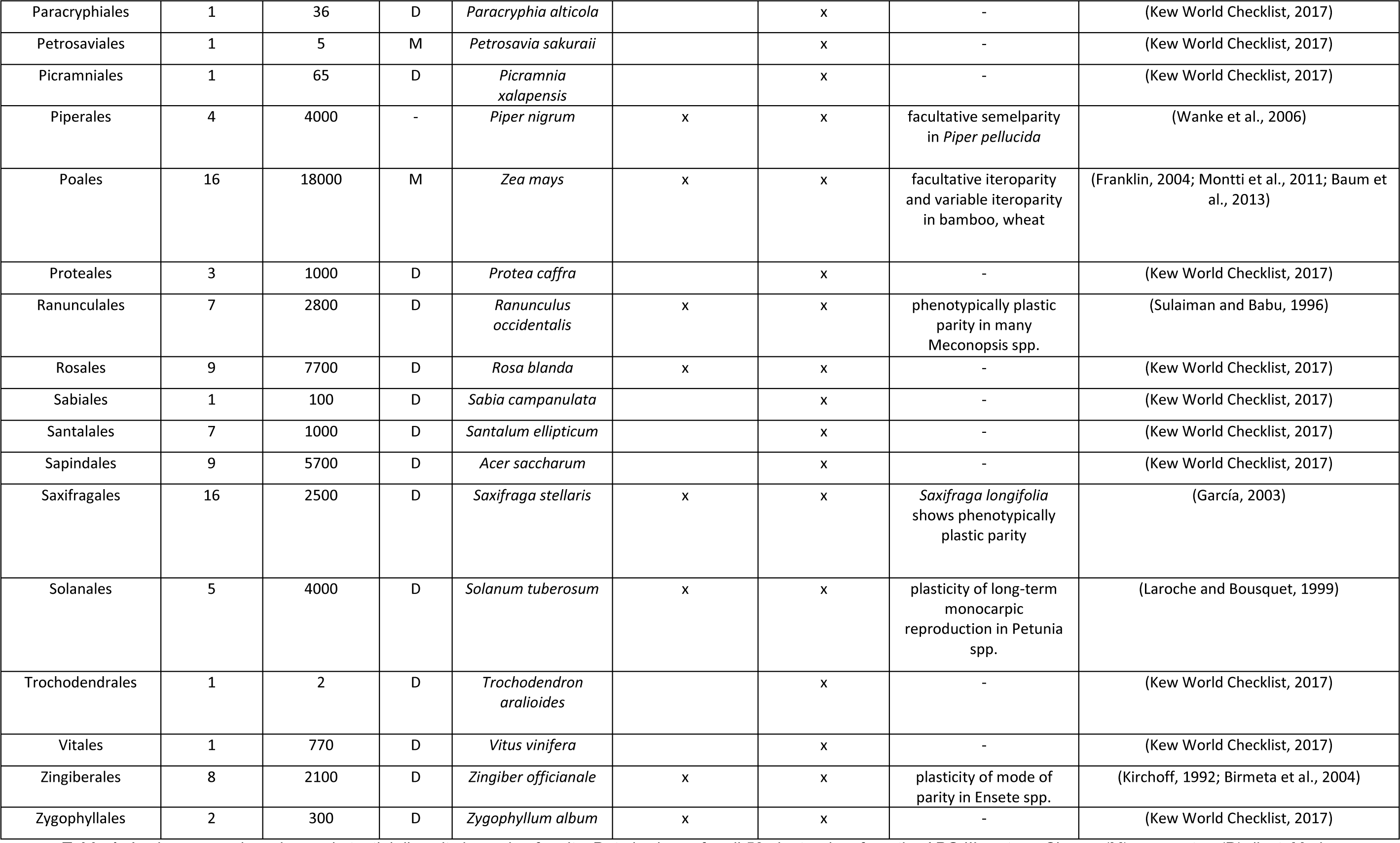
Angiosperm orders show substantial diversity in mode of parity. Data is shown for all 59 plant orders from the APG III system. Class = (M) monocot or (D) dicot. Marks indicate whether the indicated reproductive strategy is present in that order.

| Plant Order | Families | Estimated Number of Species | Class | Exemplar Species | Semelparous Species Present | Iteroparous Species Present | Notes | Reference |
| --- | --- | --- | --- | --- | --- | --- | --- | --- |
| Acorales | 1 | 2 | M | *Acorus calamus* |  | x | - | (Kew World Checklist) |
| Alismatales | 13 | 4500 | M | *Anthurium andraeanum* | x | x | facultative semelparity | (Haggard and Tiffney, 1997; Haston et al., 2009) |
| Amborellales | 1 | 1 | - | *Amborella trichopoda* | x |  | - | (Kew World Checklist, 2017) |
| Apiales | 7 | 5500 | D | *Daucus carota* | x | x | facultative semelparity in *Heracleum* spp. | (Chandler and Plunkett, 2004; Kew World Checklist, 2017) |
| Aquifoliales | 5 | 600 | D | *Ilex aquifolium* | x | x | - | (Kew World Checklist, 2017) |
| Arecales | 1 | 2600 | M | *Cocos nucifera* | x | x | facultative semelparity in *Rhopalostylis sapida* | (Silvertown et al., 1993) |
| Asparagales | 14 | 36000 | M | *Asparagus officinalis* | x | x | facultative semelparity in *Yucca whipplei* | (Keeley et al., 1986) |
| Asterales | 11 | 27500 | D | *Helianthus annuus* | x | x | facultative iteroparity in *Lobelia inflata* | (Hughes and Simons, 2014b; c) |
| Austrobaileyales | 3 | 100 | NA | *Illicium verum* |  | x | - | (Palmer et al., 2004) |
| Berberidopsidales | 2 | 6 | D | *Berberidopsis corallina* |  | x | - | (Kew World Checklist, 2017) |
| Brassicales | 17 | 4500 | D | *Brassica oleracea* | x | x | "perpetual flowering" under genetic control (*Arabis alpina*); phenotypically plastic semelparity in *Brassica campestris* | (Biswas and Mandal, 1987; Wang, Farrona, Vincent, Fornara, et al., 2009; Kew World Checklist, 2017) Albani PEP1; 2: Biswas, A. K., and S. K. Mandal. "Regulation of monocarpic senescence of *Brassica campestris* by the developing pods." *Physiologia Plantarum* 71.1 (1987): 89-94. |
| Bruniales | 2 | 80 | D | *Desfontainia spinosa* | x | x | fire-dependent facultative parity (Bruniaceae spp.) | (van Wilgen and Forsyth, 1992) Van Wilgen and Forsyth (1992) |
| Buxales | 3 | 120 | D | *Buxus sempervirens* |  | x | - | (Kew World Checklist, 2017) |
| Canellales | 2 | 136 | D | *Canella winteriana* | x | x | - | (Kew World Checklist, 2017) |
| Caryophyllales | 33 | 11000 | D | *Silene dioica* | x | x | Plasticity of iteroparous reproduction in *Ferocactus wislizeni*; variation in many long-lived Cactaceae perennial spp.; similar changes in Buckwheat (Polygonaceae) | (Bowers, 2000; Song et al., 2013) |
| Celastrales | 2 | 1300 | D | *Celastrus orbiculatus* |  | x | - | (Kew World Checklist, 2017) |
| Ceratophyllales | 1 | 10 | - | *Ceratophyllum submersum* | x | x | plasticity of iteroparous life histories in European hornworts | (Bisang et al., 2008) |
| Chloranthales | 1 | 75 | - | *Sarcandra glabra* |  | x | - | (Kew World Checklist, 2017) |
| Commelinales | 5 | 850 | M | *Commelina communis* | x | x | Phenotypically plastic semelparity in genus *Commelina*; varies from near-uniparity to extended semelparity | (Faden, 1993, 2006) |
| Cornales | 6 | 600 | D | *Hydrangea macrophylla* | x | x | Facultative iteroparity in stickleaf spp. | (Keeler, 1987) |
| Crossosomatales | 7 | 80 | D | *Crossosoma bigelovii* |  | x |  | (Kew World Checklist, 2017) |
| Cucurbitales | 8 | 2600 | D | *Begonia obliqua* | x | x | facultative iteroparity among cultivated begonias | (De Wilde, 2010) |
| Dilleniales | 1 | 400 | D | *Hibbertia stellaris* | x | x | facultative semelparity in *Hibbertia* spp | (Stebbins and Hoogland, 1976) |
| Dioscoreales | 3 | 1050 | M | *Dioscorea rotundata* |  | x | - | (Kew World Checklist, 2017) |
| Dipsacales | 2 | 1100 | D | *Lonicera periclymenum* |  | x | - | (Kew World Checklist, 2017) |
| Ericales | 25 | 11000 | D | *Vaccinium macrpocarpon* | x | x | Facultative semelparity and facultative iteroparity in *Impatiens* spp. | (Vervoort et al., 2011) |
| Escalloniales | 1 | 130 | D | *Escallonia bifida* | x | x | - | (Kew World Checklist, 2017) |
| Fabales | 4 | 20000 | D | *Pisum sativum* | x | x | facultative iteroparity in many spp. | (Nichols et al., 2007) |
| Fagales | 7 | 2000 | D | *Quercus alba* |  | x | - | (Kew World Checklist, 2017) |
| Garryales | 2 | 18 | D | *Garrya congdonii* |  | x | - | (Kew World Checklist, 2017) |
| Gentianales | 5 | 16000 | D | *Gentiana verna* | x | x | plasticity of iteroparous reproduction in long lived perennial *Frasera caroliniensis* | (Threadgill et al., 1981) |
| Geraniales | 5 | 900 | D | *Geranium rotundifolium* | x | x | - | (Kew World Checklist, 2017) |
| Gunnerales | 2 | 55 | D | *Gunnera manicata* | x | x | facultative semelparity in *Gunnera herteri* | (Wanntorp et al., 2002) |
| Huerteales | 4 | 20 | D | *Huertia cubensis* |  | x | - | (Kew World Checklist, 2017) |
| Lamiales | 20 | 24000 | D | *Lamium purpureum* | x | x | - | (Kew World Checklist, 2017) |
| Laurales | 7 | 2500 | D | *Laurus nobilis* |  | x | - | (Kew World Checklist, 2017) |
| Liliales | 10 | 1300 | M | *Lilium candidum* | x | x | Many perennial Agave spp. have substantial phenotypic plasticity in parity | (Nobel, 1977; Arizaga and Ezcurra, 1995; Rocha et al., 2005) |
| Magnoliales | 6 | 5000 | - | *Magnolia virginiana* | x | x | - | (Kew World Checklist, 2017) |
| Malpighiales | 35 | 16000 | D | *Malpighia glabra* | x | x | - | (Kew World Checklist, 2017) |
| Malvales | 9 | 6000 | D | *Malva sylvestris* | x | x | - | (Kew World Checklist, 2017) |
| Myrtales | 9 | 11000 | D | *Myrtus communis* | x | x | - | (Kew World Checklist, 2017) |
| Nymphaeales | 3 | 70 | - | *Nymphaea lotus* |  | x | - | (Kew World Checklist, 2017) |
| Oxalidales | 7 | 1800 | D | *Oxalis acetosella* | x | x | - | (Kew World Checklist, 2017) |
| Pandanales | 5 | 1300 | M | *Pandanus utilis* |  | x | - | (Kew World Checklist, 2017) |
| Paracryphiales | 1 | 36 | D | *Paracryphia alticola* |  | x | - | (Kew World Checklist, 2017) |
| Petrosaviales | 1 | 5 | M | *Petrosavia sakuraii* |  | x | - | (Kew World Checklist, 2017) |
| Picramniales | 1 | 65 | D | *Picramnia xalapensis* |  | x | - | (Kew World Checklist, 2017) |
| Piperales | 4 | 4000 | - | *Piper nigrum* | x | x | facultative semelparity<br>in *Piper pellucida* | (Wanke et al., 2006) |
| Poales | 16 | 18000 | M | *Zea mays* | x | x | facultative iteroparity<br>and variable iteroparity<br>in bamboo, wheat | (Franklin, 2004; Montti et al., 2011; Baum et al., 2013) |
| Proteales | 3 | 1000 | D | *Protea caffra* |  | x | - | (Kew World Checklist, 2017) |
| Ranunculales | 7 | 2800 | D | *Ranunculus occidentalis* | x | x | phenotypically plastic<br>parity in many<br>*Meconopsis* spp. | (Sulaiman and Babu, 1996) |
| Rosales | 9 | 7700 | D | *Rosa blanda* | x | x | - | (Kew World Checklist, 2017) |
| Sabiales | 1 | 100 | D | *Sabia campanulata* |  | x | - | (Kew World Checklist, 2017) |
| Santalales | 7 | 1000 | D | *Santalum ellipticum* |  | x | - | (Kew World Checklist, 2017) |
| Sapindales | 9 | 5700 | D | *Acer saccharum* |  | x | - | (Kew World Checklist, 2017) |
| Saxifragales | 16 | 2500 | D | *Saxifraga stellaris* | x | x | *Saxifraga longifolia*<br>shows phenotypically<br>plastic parity | (García, 2003) |
| Solanales | 5 | 4000 | D | *Solanum tuberosum* | x | x | plasticity of long-term<br>monocarpic<br>reproduction in *Petunia*<br>spp. | (Laroche and Bousquet, 1999) |
| Trochodendrales | 1 | 2 | D | *Trochodendron aralioides* |  | x | - | (Kew World Checklist, 2017) |
| Vitales | 1 | 770 | D | *Vitis vinifera* |  | x | - | (Kew World Checklist, 2017) |
| Zingiberales | 8 | 2100 | D | *Zingiber officinale* | x | x | plasticity of mode of<br>parity in *Ensete* spp. | (Kirchoff, 1992; Birmeta et al., 2004) |
| Zygophyllales | 2 | 300 | D | *Zygophyllum album* | x | x | - | (Kew World Checklist, 2017) |

In this review I begin by reviewing the development of both the discrete and continuous conceptions of parity as evolutionary hypotheses and/or models. Next, I review empirical work that highlights the existence of natural variation in reproduction along a semelparity-iteroparity continuum, focusing on three distinct strategies: facultative iteroparity, facultative semelparity, and phenotypically plastic parity. I conclude by exploring the implications of the continuous conception of parity for: (1) the study of seasonality as a “digitization” of reproduction, (2) the process of mathematically modelling life-history optimization, and (3) the study of the molecular regulation of reproductive traits linked to parity.

## II. The Discrete Conception of Parity

### (1) ‘Cole’s Paradox’ and the development of the discrete conception of parity

Although the first mathematical model of the intrinsic rate of increase of annual plants was constructed by Linnaeus (1744), Lamont Cole (1954) was the first to categorize life histories into dichotomous “semelparous” and “iteroparous” groups: a semelparous organism is one that “dies upon producing seed” and therefore “potential population growth may be considered on the assumption that generations do not overlap” (p. 109), while iteroparous organisms include a variety of cases, from those where “only two or three litters of young are produced in a lifetime” as well as “various trees and tapeworms, where a single individual may produce thousands of litters” (p. 118). Thus, Cole created, and contemporary theorists have inherited, a conception of parity as a discrete variable: an organism either reproduces more than once or it doesn’t.

Cole also identified “the paradox of semelparity”, and wrote that “for an annual species, the absolute gain in intrinsic population growth which could be achieved by changing to the perennial reproductive habit would be exactly equivalent to adding one individual to the average litter size” (Cole 1954, p. 118). Consequently, according to the model he developed, a semelparous or iteroparous strategy evolves in response to strong directional selection for trait-values that: (1) maximize the annual rate of intrinsic increase; and (2) are subject to tradeoffs, since reproductive effort is always limited by resource availability. The “paradox of semelparity” is that the relative intrinsic rates of increase for semelparous and iteroparous strategies are very similar (i.e. they differ only by one individual—the mother), which suggests that iteroparity, not semelparity, should be rare, while in nature, iteroparous life histories are generally more common than semelparous ones. Cole’s articulation of the paradox of semelparity motivated many studies searching for theoretical selective advantages of traits linked to discrete semelparous and iteroparous strategies (Murdoch, 1966; Murphy, 1968; Omielan, 1991; Su and Peterman, 2012; Vaupel et al., 2013; Cushing, 2015), as well as attempts to detect these selective advantages in natural systems (Murphy and Rodhouse, 1999; Kaitala et al., 2002; Kraaijeveld et al., 2003; Franklin and Hogarth, 2008; Gagnon and Platt, 2008; Fisher and Blomberg, 2011). Following Cole, semelparous strategies considered in later life history models were usually also annual (although empirical data on long-lived semelparous organisms was also collected, e.g. Young and Augspurger, 1991; García, 2003), and thus the primary goal of many models purporting to explain the evolution of semelparity was to provide reasons why a perennial-iteroparous strategy might confer higher fitness than an annual-semelparous one.

Cole’s “paradox of semelparity” was resolved by acknowledging that differences in age-specific rates of mortality affect the relative fitness of semelparous and iteroparous habits. Building on prior analytical work (see: Murphy, 1968; Emlen, 1970; Gadgil and Bossert, 1970; Bryant, 1971), Charnov and Schaffer (1973) and Schaffer (1974b) noted that the expected fitness value of individuals at juvenile (i.e. prereproductive) and adult (i.e. reproductively mature) developmental stages often differed. They then argued that when the survival of adults was more assured than the survival of juveniles, an iteroparous habit would have a growth advantage over a semelparous one. Thus, their model emphasized that the value of the age class with a lower age-or stage-specific rate of mortality was—assuming equal fitness across age classes—greater than the value of the age class with a higher rate of mortality. Young (1981) extended this insight into a more general model of intrinsic rates of increase, which incorporated not only differences in age-specific survivorship, but also differences in prereproductive development time and time between reproductive episodes. This model provided three major reasons why semelparity might be favoured by natural selection. First, high adult mortality—or the early onset of reproductive senescence—might prevent iteroparous species from accruing fitness gains from established parents over long timescales. Second, a high population growth rate should favour semelparity outright. Third, when the marginal cost of additional offspring is inversely proportional to the number of offspring produced, fecundity is maximized by investing all reproductive effort into a single episode, i.e. adopting an extreme annual-semelparous life history—see also Schaffer (1974) and Schaffer and Gadgil (1975).

### (2) Further theoretical work on parity as a discrete trait

Given that earlier work sought to explain the prevalence of semelparous and iteroparous strategies by identifying differences in age-specific mortality, recent work has sought to explain why differences in age-specific mortality persist, as well as how varying environmental conditions facilitate the co-existence of different modes of parity. Bulmer (1985, 1994) incorporated a model that incorporated density-dependence, a model that was generalized by Ranta et al. (2002). Another common approach has been to use simulations, based on comparisons between discrete strategies, to argue that spatial heterogeneity and stochastic events (i.e. demographic disasters and windfalls) influence the evolutionary stability of each mode of parity over small spatial scales (e.g. Ranta et al., 2001). Similarly, Zeineddine and Jansen (2009) examined the role that discrete modes of parity may play in evolutionary tracking, suggesting that species adopting an annual-semelparous strategy may have an evolvability advantage over perennial-iteroparous congeners. Moreover, considerable evidence now supports two general conclusions: (1) that theoretically optimal life history strategies strongly depend on optimizing parity (Maltby and Calow, 1986; Keeley and Bond, 1999; Iguchi and Tsukamoto, 2001; Stegmann and Linsenmair, 2002; Kraaijeveld et al., 2003; Leiner et al., 2008; Trumbo, 2013), but also (2) that parity is especially important for predicting reproductive scheduling (Schaffer and Gadgil, 1975; Kozłowski and Wiegert, 1986; Iwasa, 1991; Kozłowski, 1992; McNamara, 1997; Cooke et al., 2004; Miller et al., 2012; Vaupel et al., 2013; Oizumi, 2014), and/or the optimal allocation of reproductive effort to offspring (Cohen, 1966; Smith and Fretwell, 1974; Winkler and M., 2002; Einum and Fleming, 2007; Gremer and Venable, 2014; Mironchenko and Kozłowski, 2014). Thus the current theoretical life history literature is replete with papers discussing the knock-on effects of the assumption that modes of parity are discrete rather than continuous.

### (3) Empirical support for the discrete conception of parity

Empirical support for the discrete model of parity is strongest where perennial-iteroparous and annual-semelparous (or, rarely, perennial-semelparous) congeneric species have starkly different life histories. For instance, in a comparison of Mount Kenya species of the genus *Lobelia*, Young (1984) found that juvenile and adult mortality of the annual-semelparous species *L. telekii* were higher than in the closely related perennial-iteroparous species *Lobelia keniensis*. Young concluded that the difference in age-specific rates of mortality would strongly influence the expected value of future reproduction for each species, leading to perennial-iteroparity in one species and annual-semelparity in the other (Young, 1990). Similar comparisons between semelparous and iteroparous congeners or confamilials have been conducted in insects (Fritz et al., 1982; Stegmann and Linsenmair, 2002), salmon (Dickhoff, 1989; Unwin et al., 1999; Crespi and Teo, 2002; Kindsvater et al., 2016), snakes (Bonnet, 2011), algae (De Wreede and Klinger, 1988) and dasyurid marsupials (Kraaijeveld et al., 2003; Mills et al., 2012).

Other studies have focused on reproductive effort, since a declining marginal cost of offspring in terms of reproductive effort should select for an annual-or perennial-semelparous life history over an perennial-iteroparous one. This is the cited cause of the evolution of semelparity in *Digitalis purpurea* (Sletvold, 2002), and in *Antechinus agilis* (Smith and Charnov, 2001; Fisher and Blomberg, 2011). Growth rate is also important; in two subspecies of *Yucca whipplei*, the semelparous variant showed higher viability and faster time to germination than the iteroparous variant did (Huxman and Loik, 1997). Further studies highlight the mortality differences between juveniles and adults, which explains the evolution of semelparity in a variety of long-lived semelparous plants (Foster, 1977; Kitajima and Augspurger, 1989; Young and Augspurger, 1991), as well as in salmonids (Fleming, 1998; Crespi and Teo, 2002; Hendry et al., 2004; Sloat et al., 2014).

Models have been developed that use demographic parameters to predict age and size at first flowering for semelparous (monocarpic) plants; however, these models have been found to be more appropriate for long-lived than short-lived species (Metcalf et al 2003, Rees and Rose 2002). More recently, mathematical modelling of evolutionary responses to discrete semelparous and iteroparous strategies has focused on whether the maintenance of both modes of parity can be a consequence of stochasticity in the ratio of juvenile to adult mortality (Murphy, 1968; Ranta et al., 2002), of differences in the effects of density on age-specific mortality (Bulmer, 1985), or as a consequence of population instability (Ranta et al., 2000).

## III. The continuous conception of parity

### (1) From uniparity to continuous reproduction

However, in many cases substantial unexplained variation in parity exists even after factors such as age-specific mortality, density dependence, and environmental effects are taken into account. For this reason it seems as though models based on the discrete conception of parity describe a limited range of special cases and not the majority of systems with congeneric or confamilial species with different reproductive strategies. This problem arises because theoretical models of the discrete conception of parity make two characteristic assumptions. First, these models assume that reproductive output is allocated among cycles (typically seasons or years) rather than expressed continuously. This means that offspring produced at two different times within a single season are “counted” as being part of the same reproductive episode, while offspring produced at two different times in two different seasons are counted as part of categorically different reproductive episodes. This permits the calculation of threshold values (e.g. of size or age) beyond which selection should begin to favour one mode of parity or the other, but is based on a distinction that is arbitrary. Second, each individual is assumed to express a single reproductive strategy; models do not predict phenotypically plastic modes of parity, or facultative switching between modes.

These assumptions do not hold in many cases. The fact that semelparous reproduction rarely occurs “once”—i.e. in exactly one place, at exactly one time—has led to a new treatment of parity as continuously varying between extremes of “pure” semelparous and iteroparous reproduction. This approach has gained traction because there is considerable ambiguity in “breeding once” (Kirkendall and Stenseth, 1985). Moreover, “annuality” and “perenniality”—terms that refer to the number of years in which organisms reproduce—cannot be used interchangeably with “semelparity” and “iteroparity”, which refer to the number of reproductive episodes organisms have (Fritz et al., 1982; Kirkendall and Stenseth, 1985). In “The Evolution of Life Histories”, Roff (1992) noted that, “if we consider our unit of time to be a single year, annuals can be termed semelparous and perennials iteroparous. A further division is possible within annuals, for some reproduce once and are, therefore, semelparous within any time scale, while others flower repeatedly throughout the summer and, hence, are iteroparous with respect to annuals that flower only once, but semelparous with respect to perennials” (p. 248). That is, it is the simultaneity and the finality of the reproductive episode (i.e. the concentration of reproductive effort) that defines perfect semelparity. Therefore, the continuous conception characterizes “extreme” semelparity to be a single, complete, and exhaustive reproductive episode where all reproductive effort is invested at once. Examples of this strategy—which Kirkendall and Stenseth (1985) termed “uniparity”— include mayflies and mites of the genus Adactylidium (Edmunds et al., 1976; Corkum et al., 1997). Both male and female mayflies die shortly after mating and dispersing fertilized eggs. In Adactylid mites, offspring devour the mother from the inside out, and are thus obligately annual-semelparous (Elbadry and Tawfik, 1966; Goldrazena et al., 1997). The correspondingly “extreme” perennial-iteroparous strategy is a long-lived perennial strategy that spreads reproductive effort out evenly among a very large number of reproductive cycles. Many species, including bristlecone pine, many deep-sea zoanthids, and other supercentennial species that reproduce regularly show such a habit (Baker, 1992; Druffel et al., 1995; Finch, 1998; Rozas, 2003). Intermediate strategies complete reproduction over a shorter timescale than bristlecone pine, but over a longer timescale than Adactylid mites.

The continuous conception of parity is therefore very simple: parity is a trait like any other, and continuous variation in factors affecting reproduction (e.g. resource availability, timing of reproduction, etc.), where subject to evolution by natural selection, should result in continuous variation in reproductive strategy (see: Salguero-Gómez et al., 2016). Furthermore, rather than considering only whether organisms complete reproduction within a given year, and making no finer distinction, life history strategies should be compared by the degree to which they concentrate or disperse reproductive effort—and hence risk of reproductive failure—in time. For example, a mature biennial strategy (where an organism reproduces once per year in two consecutive years) distributes reproductive effort over a shorter timescale than does a long-lived perennial congener (where an organism reproduces once per year in many years); although the biennial strategy is not semelparous, it is further toward the “uniparous” end of the continuum of modes of parity than is the perennial strategy. Similarly, an annual-semelparous life history that reproduces rapidly lies further toward this end of the continuum than does an annual-semelparous life history in which reproduction is spread over a longer period of time.

### (2) Empirical support for the continuous conception of parity

There is considerable empirical support, from lab and field studies alike, for the notion that parity varies continuously. These results have made it increasingly clear that demographic explanations rooted in the discrete model of parity fail to provide an universal evolutionary explanation for the appearance of different modes of parity. The most obvious objections to the universal applicability of this theory come from species that are facultatively semelparous, from species that reproduce irregularly or opportunistically, or from comparisons between related iteroparous and semelparous species that do not show measurable differences in factors affecting intrinsic rates of increase, including age-specific rates of mortality. These situations are not uncommon in nature. The problem they present is significant because the evolutionary transition from semelparity to iteroparity (and back) is ubiquitous, and has occurred in a wide variety of taxa (see Table 1 for an example using data from angiosperm orders).

There are important consequences for adopting the continuous conception of parity as a starting point for modelling the evolution of different modes of parity. Mathematical models based on the discrete conception of parity often predict threshold values—in mortality rate, size at initiation of reproduction, or expected growth rate—that do not agree with empirical observation (Omielan, 1991; Lessells, 2005; Piñol and Banzon, 2011; Su and Peterman, 2012; Trumbo, 2013; Vaupel et al., 2013). In particular, ESS models derived from assumptions rooted in the discrete conception of parity frequently underestimate the adaptive value of semelparous reproductive strategies; even after accounting for the effects of environmental stochasticity and density-dependence, ESS models predict that semelparous strategies should be less abundant—and less fit— than they have been found to be (Benton and Grant, 1999). In addition, there are empirical cases that explicitly do not conform to the predictions of the discrete model. For example, an analysis of 12 winter-establishing primrose species (Oenothera: Onagraceae) found no significant differences in mortality estimates or in environmental determinants of fitness for semelparous and iteroparous species (Evans et al., 2005). In some cases, the problem may be that life histories are too complex for organisms to follow discrete strategies; many salmon species also do not fit neatly into “classical” annual-or perennial-semelparous and perennial-iteroparous classifications (Unwin et al., 1999; Hendry et al., 2004). Other research has suggested that deterministic models of investment may better fit long-lived than short-lived semelparous species, given that many annual semelparous species (usually plants) show substantial phenotypic plasticity in phenology (e.g. size at first flowering), offspring quality and overall fecundity (Burd et al., 2006).

In order to provide a coherent exposition of the extensive body of recent work that shows empirical support for the continuous conception of parity, in what follows I focus on three “intermediate” life histories that are neither annual-semelparous nor perennial-iteroparous, but express another mode of parity that falls somewhere in between. These include: (1) facultative iteroparity; (2) facultative semelparity; and (3) phenotypically plastic parity. Although these three examples are the most common modes of parity that are neither classically (annual-or perennial-) semelparous nor iteroparous, other intermediate strategies exist, particularly when species or clades have idiosyncratic life histories. In addition, an individual population or species may itself express multiple modes of parity. For instance, arctic cod (*Boreogadus saida*) are annual-semelparous in nature, but can reproduce in two consecutive years in captivity, making them facultatively iteroparous (Hop et al., 1995; Hop and Gjosaeter, 2013). However, males and females of this species also seem to have different life histories – males begin to reproduce at an earlier age, and can allocate extreme amounts of reproductive effort to a single instance of reproductive activity; semelparity in this species is thus also phenotypically plastic (Nahrgang et al., 2014). Although examples of each life history are provided below, many more have been added to Table 2, a list of species showing facultatively varying or plastic modes of parity.

**Table 2.** Species known to display facultative semelparity, facultative iteroparity, a continuum of modes of parity, or phenotypic plasticity with respect to mode of parity.

| Focal Species | Clade | Continuously varying traits identified | Facultative Iteroparity | Facultative Semelparity | Continuous Variation in Parity | Phenotypically Plastic Parity | Reference |
| --- | --- | --- | --- | --- | --- | --- | --- |
| *Acer negundo* | Angiosperm | timing of reproduction |  |  |  | x | (Lamarque et al., 2015) |
| *Agave celsii*, *Agave difformis* | Angiosperm | timing of reproduction;<br>reproductive effort |  |  | x | x | (Rocha et al., 2005) |
| *Allogamus uncatus* | Insect | timing of reproduction;<br>duration of reproduction |  |  | x | x | (Shama and Robinson, 2009) |
| *Alosa sapidissima* | Fish | timing of reproduction; clutch size |  | x | x | x | (Leggett and Carscadden, 1978) |
| *Amblyrhynchus cristatus* | Reptile | timing of reproduction |  |  | x | x | (Vitousek et al., 2010) |
| *Antechinus stuartii* | Mammal | adult mortality | x |  | x |  | (Fisher and Blomberg, 2011) |
| *Arabidopsis lyrata* | Angiosperm | timing of reproduction;<br>duration of reproduction |  |  | x |  | (Remington et al., 2015) |
| *Arabis fecunda* | Angiosperm | reproductive effort |  |  | x | x | (Lesica and Shelly, 1995) |
| *Bambusa narnhemica* | Angiosperm | timing of reproduction |  |  | x |  | (Franklin, 2004) |
| *Beta vulgaris* | Angiosperm | reproductive effort |  |  |  | x | (Letschert, 1993; Hautekèete et al., 2001, 2002, 2009) |
| *Boreogadus saida* | Fish | adult mortality;<br>reproductive effort |  |  | x |  | (Nahrgang et al., 2014) |
| *Botryllus sclosseri* | Tunicate | timing of reproduction, clutch size | x |  | x | x | (Grosberg, 1988; Harvell and Grosberg, 1988) |
| *Bradybaena pellucida* | Gastropod | duration of reproduction |  |  | x | x | (Nyumura and Asami, 2015) |
| *Cynodonichthys brunneus*, *Cynodonichthys magdalenae*, *Cynodonichthys kuelpmanni*, *Anablepsoides immaculatus*, and *Laimosemion frenatus* | Fish | timing of reproduction; diapause length |  | x | x | x | (Varela-Lasheras and Van Dooren, 2014) |
| *Daphnia galeata* | Crustacean | timing of reproduction |  |  | x | x | (Henning-Lucass et al., 2016) |
| *Daucus carota* | Angiosperm | timing of reproduction |  |  | x | x | (Lacey, 1988) |
| *Digitaria californica* | Angiosperm | juvenile mortality |  |  | x |  | (Smith et al., 2000) |
| *Dosidicus gigas* | Mollusc | timing of reproduction; duration of reproduction | x |  | x | x | (Hoving et al., 2013) |
| *Emoia atroscostata* | Reptile | timing of reproduction |  |  | x |  | (Alcala and Brown, 1967) |
| *Erpobella octoculata* | Leech | adult mortality; reproductive effort | x |  | x | x | (Maltby and Calow, 1986) |
| *Erysimum capitatum* | Angiosperm | timing of reproduction |  | x | x |  | (Kim and Donohue, 2011) |
| *Eulamprus tympanum* | Reptile | clutch size; timing of reproduction |  |  | x | x | (Doughty and Shine, 1997) |
| *Forficula auricularia* | Insect | timing of reproduction |  |  |  | x | (Meunier et al., 2012) |
| *Fucus serratus*; *Himanthalia elongata* | Algae | duration of reproduction |  |  | x |  | (Brenchley et al., 1996) |
| *Gaimardia bahamondei* | Bivalve | timing of reproduction; duration of reproduction | x |  | x | x | (Chaparro et al., 2011) |
| *Galaxias maculatus* | Fish | timing of reproduction | x |  |  | x | (Stevens et al., 2016) |
| *Gasterosteus aculeatus* | Stickleback | timing of reproduction; reproductive effort |  | x | x | x | (Snyder, 1991; Bell and Foster, 1994; Baker et al., 2008, 2015) |
| *Gracilinanus microtarsus* | Mammal | male adult mortality | x |  | x | x | (Kraaijeveld et al., 2003; Martins et al., 2006) |
| *Idiosepius pygmaeus* | Cephalopod | timing of reproduction | x |  | x |  | (Lewis and Choat, 1993; Nesis, 1996) |
| *Ligia cinerascens* | Crustacean | adult mortality; timing of reproduction | x | x | x | x | (Furota and Ito, 1999) |
| *Lobelia inflata* | Angiosperm | timing of reproduction; duration of reproduction | x |  | x | x | (Hughes and Simons, 2014 a,b,c) |
| *Loligo vulgaris* | Cephalopod | timing of reproduction; clutch size | x |  | x |  | (Melo and Sauer, 1999; Sauer et al., 1999) |
| *Mallotus villosus* | Fish | adult mortality; timing of reproduction |  | x | x | x | (Christiansen et al., 2008) |
| *Marmosops paulensis* | Mammal | female adult mortality | x |  | x | x | (Leiner et al., 2008) |
| *Mimulus guttatus* | Angiosperm | duration of flowering |  |  | x | x | (Van Kleunen, 2007) |
| *Misumena vatia* | Arachnid | juvenile mortality |  |  | x | x | (Morse, 1994; Morse and Stephens, 1996) |
| *Nautilus* spp. | Cephalopod | timing of reproduction; timing of senescence | x |  | x | x | (Ward, 1983, 1987) |
| *Oenothera deltoides*; *Oenothera pallida* | Angiosperm | timing of reproduction; adult mortality | x | x | x |  | (Evans et al., 2005) |
| *Oncorhynchus mykiss* | Fish | reproductive effort | x |  | x | x | (Seamons and Quinn, 2010) |
| *Oncorhynchus nerka* | Fish | timing of reproduction |  |  |  | x | (Hendry et al., 2004) |
| *Oncorhynchus tshawytscha* | Fish | adult mortality | x |  |  | x | (Unwin et al., 1999) |
| *Onopordum illyricum* | Angiosperm | timing of reproduction |  | x |  |  | (Rees et al., 1999) |
| *Opisthoteuthis agassizii*, *Opisthoteuthis grimaldii* and *Grimpoteuthis glacialis* | Cephalopod | timing of reproduction; reproductive effort |  |  | x |  | (Aldred et al., 1983; Villanueva, 1992; Vecchione et al., 1998) |
| *Panicum bisulcatum*, *Cyperus michelianus*, *Fimbristylis miliacea*, and *Eclipta prostrata* | Angiosperm | timing of reproduction; duration of reproduction |  |  | x | x | (Song et al., 2015) |
| *Parantechnius apicalis* | Mammal | adult mortality |  | x |  | x | (Wolfe et al., 2004) |
| *Plecoglossus altivelis* | Fish | timing of reproduction; reproductive effort | x |  | x | x | (Iguchi, 1996; Iguchi and Tsukamoto, 2001) |
| *Puya raimondii* | Angiosperm | reproductive effort |  | x | x |  | (Jabaily and Sytsma, 2013) |
| *Rana arvalis* | Frog | timing of reproduction |  |  |  | x | (Richter-Boix et al., 2014) |
| *Sasa senanensis*, *Sasa kurilensis*, and *Sasa palmata* | Angiosperm | timing of reproduction; reproductive effort | x | x |  | x | (Mizuki et al., 2014) |
| *Sepia officinalis* | Cephalopod | adult mortality; timing of reproduction; clutch size; timing of senescence | x |  | x | x | (Boletzky, 1988; Rocha et al., 2001) |
| *Stegodyphus lineatus* | Arachnid | adult mortality; timing of reproduction |  |  | x | x | (Schneider and Lubin, 1997) |
| *Sthenoteuthis oualaniensis* | Cephalopod | timing of reproduction | x |  |  |  | (Rocha et al., 2001) |
| *Strix uralensis* | Bird | timing of reproduction |  |  |  | x | (Brommer et al., 2012) |
| *Uta stansburiana* | Reptile | duration of reproduction, |  | x | x | x | (Tinkle, 1969) |
| *Verbascum thapsis* | Angiosperm | timing of reproduction |  |  | x |  | (Reinartz, 1984) |
| *Wyeomyia smithii* | Angiosperm | timing of reproduction |  |  | x | x | (Bradshaw, 1986) |
| *Xerolenta obvia* | Gastropod | timing of reproduction | x |  | x | x | (Lazaridou and Chatziioannou, 2005) |
| *Xeropicta derbentina* | Gastropod | timing of reproduction | x |  | x | x | (Aubry et al., 2005; Kiss et al., 2005) |
| *Yucca whipplei* | Angiosperm | reproductive effort |  | x | x |  | (Huxman and Loik, 1997) |

#### (a) Facultative Iteroparity

Many semelparous species have shown the ability to facultatively reproduce one or more times after an initial bout of reproduction has began and ended—this is termed “facultative iteroparity”. Facultative iteroparity can be adaptive when it either: (1) provides an opportunity to realize fitness gains from an unexpected abundance of resources, or (2) shifts reproductive effort from inopportune to opportune times. The first type of adaptive facultative iteroparity occurs when additional bouts of reproduction increase fitness by permitting unexpected “bonus” resources to be invested in new offspring. For example, mothers of the semelparous crab spider *Misumena vatia* (Araeae, Thomsidae) typically lay and provision a single brood of eggs (Gertsch, 1939; Morse, 1979), however in response to high food availability and/or usually warm environmental conditions, they are capable of laying and caring for a second brood if sperm supplies are not depleted (Morse, 1994). A similar facultative double-broodedness in response to unusually favourable environment has been observed in the green lynx spider *Peucetia viridans* (Fink, 1986). In addition, a small proportion of Chinook salmon (*Onchorhynchus tshawytscha*), which typically reproduce only once, have been found to survive and reproduce in two or three additional seasons (Unwin et al., 1999). Tallamy and Brown (1999) showed that large, well-provisioned female burying beetles in multiple species in the genus Nicrophorus can reproduce more than once, despite the fact that small females can typically breed only once.

The second form of adaptive facultative iteroparity occurs when deferral of reproductive effort—from a primary reproductive episode to a secondary one—allows an organism to reproduce at a more opportune time. Reproduction is deferred to seek the highest marginal fitness return on invested reproductive effort. For example, when high organic pollution levels disrupt primary reproduction in the freshwater leech *Erpobdella octoculata*, reproduction ceases and remaining reproductive effort is deferred to a second reproductive bout produced the next year (Maltby and Calow, 1986). Similar behaviour has been seen in another Erpobdellid leech, *Erpobdella obscura* (Peterson, 1983; Davies and Dratnal, 1996) as well as in many cephalopods (Rocha et al., 2001). Adaptive deferral of reproductive effort is common in crab spiders. In *Lysiteles coronatus*, artificial brood reductions resulted in the production of a second brood, and the degree of deferral was proportional to the degree of the original reduction (Futami and Akimoto, 2005). This was also observed in the field in Eresid spiders of the genera Anelosimus and Stegodyphus, both of which facultatively produce a second brood in response to nest predation (Schneider and Lubin, 1997; Schneider et al., 2003; Grinsted et al., 2014). Although the adaptive potential of facultative iteroparity is often apparent, facultative iteroparity may also be vestigial instead of adaptive. In this case, the organism’s life history merely reflects an ancestral state, and the second (or additional) bout of reproduction should confer little or no adaptive value (Golding and Yuwono, 1994; Hughes and Simons, 2014b).

#### (b) Facultative Semelparity

Facultative semelparity occurs when species that are normally perennial-iteroparous—i.e. have multiple, discontinuous reproductive episodes that span more than one year—are capable of expressing only a single reproductive bout (Christiansen et al., 2008). This is a useful strategy for organisms to use to take advantage of unusually good environmental conditions for reproduction. For example, in the short-lived mustard *Boechera fecunda* (also known as *Arabis fecunda*; Brassicaeae), plants are capable of wide range of reproductive strategies, from near-instantaneous semelparity to multi-year iteroparity. This is because *B. fecunda* can produce many small axillary inflorescences in any given year, and their production does not preclude flowering by the same rosette in the subsequent year. However, plants can also produce large “terminal inflorescences” that exhaust remaining resources and lead to senescence and death. Although some plants produce axillary inflorescences for several years before a terminal inflorescence, others produce a terminal inflorescence in their first year (Lesica and Shelly, 1995; Lesica and Young, 2005). A similar system is seen in common foxglove, *Digitalis purpurea* (Scrophulariaceae), which is predominantly biennial or perennial-iteroparous, but can be facultatively semelparous if resource availability in the first year is high (Sletvold, 2002). Facultative semelparity has also been observed in capelin (Christiansen et al., 2008; Loïc et al., 2012), squid, soil microarthropods (Siepel, 1994), dasyurid marsupials (Kraaijeveld et al., 2003; Martins et al., 2006), and in the flowering plants *Ipomopsis aggregata* (Silvertown and Gordon, 1989) and *Cynoglossum officinale* (Williams, 2009). Some facultatively semelparous species show a continuous range of types of reproductive episode, rather than discretely fatal or non-fatal ones. *Erysimum capitatum* (Brassicaceae) produces multiple reproductive episodes in environments where water is plentiful; however, where water is scarce it expresses a semelparous strategy (Kim and Donohue, 2011).

#### (c) Phenotypically Plastic Parity

Like other traits, mode of parity can be phenotypically plastic. This occurs when different modes of parity are expressed by a single species in response to environmental cues. Different modes of parity can occur both: (1) among individuals; or (2) within the reproductive episode of a single individual. Phenotypically plastic mode of parity is common source of intraspecific variation in life history characters; thus this is a major source of confusion for mathematical models that predict a single optimal value for all semelparous and all iteroparous habits. Such differences need not be dramatic—for example, as a consistent response to environmental triggers, even small differences in the length of a semelparous reproductive episode may have significant effects on fitness.

Strong empirical evidence of phenotypically plastic mode of parity is found in sea beets (Beta spp., Amaranthaceae), which display reproductive strategies along “a gradient from pronounced iteroparity to pronounced semelparity” (Hautekèete et al., 2001, p. 796). Interestingly, the selective pressures faced by these species seem to elicit similar plastic responses. Environmental stressors cause individuals to trade off future fecundity for increased immediate reproductive effort, resulting in a parity gradient tending to semelparity when environmental stress becomes intense (Hautekèete et al., 2001, 2009). This pattern is consistent with the prediction that higher current reproductive effort can prevent organisms from being exposed to uncertain or risky environments (Williams, 1966; Vahl, 1981; Rubenstein, 2011; Trumbo, 2013). Similar trade-offs were observed in *Yucca whipplei* (Huxman and Loik, 1997), *Chusquea ramosissima* (Montti et al., 2011), and *Onopordum illyricum* (Rees et al., 1999). *Lobelia inflata* is capable of producing different semelparous strategies, from a nearly instantaneous annual-semelparity, where plants produce many similar flowers quickly and simultaneously, to (nonadaptive) facultative biennial-iteroparity, where as much as half of all reproductive effort is invested in a second reproductive episode. The time of initiation of reproduction strongly predicted which of these strategies is realized (Hughes and Simons, 2014b; c).

Many insect species are also capable of displaying a range of modes of parity among individuals (Trumbo, 2013). In the assassin bug (*Atopozelus pallens*), females deposit eggs in small clutches, approximately every two days. However, the number of clutches—and hence how prolonged this reproductive episode is—varies substantially (Tallamy et al., 2004). Similarly, European earwigs (*Forficula auricularia*) show condition-dependent semelparity; females either deposit all eggs into a single clutch or lay two clutches (Meunier et al., 2012; Ratz et al., 2016). Most insects showing variation in the number of clutches produced do so in response to abiotic cues, particularly temperature and day length (Bradshaw, 1986). This behavior can also be found in ascidarians (Grosberg, 1988) and semelparous mammals (Wolfe et al., 2004; Mills et al., 2012).

Phenotypic plasticity within a reproductive episode of a single individual is noticeable when a semelparous organism displays a changing reproductive strategy— varying along the continuum of parity—that cannot be attributed to developmental, environmental, or architectural constraints (Diggle, 1995, 1997). This pattern is more difficult to detect than phenotypically plastic strategies that differ between individuals, but in many systems observable differences exist between the “packaging” of reproductive effort, resulting in adaptive variation in phenology or offspring quality through time. This can be difficult, since because they reproduce only once, semelparous organisms are expected to show high reproductive effort (Bonser and Aarssen, 2006). However, the development of fruits of the semelparous plant *Lobelia inflata* varied continuously; in this system, late fruits showed accelerated phenology and higher offspring quality relative to early fruits. This pattern, which indicated that more reproductive effort was invested in later fruit, shows that *L. inflata* does not “reproduce once” but dynamically allocates reproductive effort throughout a sequence of repeated fruiting events ((Hughes and Simons, 2014a, 2015). Likewise, in populations of the semelparous plant *Centaurea corymbosa*, plants showed highly variable life cycles— dynamically varying the proportion of reproductive effort allocated to sequential flowers—depending on environmental conditions and crowding (Acker et al., 2014).

### (3) Evolutionary transitions between modes of parity are ubiquitous

Transitions between different strategies along the semelparity-iteroparity continuum are common throughout the tree of life. Furthermore, modes of parity appear to be evolutionarily labile within species, and many species show significant intraspecific differences in the expression of parity (e.g. Maltby and Calow, 1986; Hughes and Simons, 2014a; c). Thus while some clades consistently display a single mode of parity (e.g. most placental mammals are exclusively iteroparous), many clades show considerable variability (see Table 1 for data from plant orders). Among cephalopoda (Mollusca), *Loligo opalescens*, *Octopus vulgaris*, *O. mimus*, and *O. cyanea* display extreme semelparity (Ikeda et al., 1993; Rocha et al., 2001), while *Nautilus* spp. show extreme perennial-iteroparity (Ward, 1983, 1987). However, other cephalopods, including *Octopus chierchiae*, *Sthenoteuthis oualaniensis*, and *Dosidicus gigas* show varying degrees of facultative iteroparity (Nesis, 1996; Laptikhovsky, 1998, 1999; Rocha et al., 2001), while still others, including *Sepia officinalis*, *Loligo vulgaris*, *L. bleekeri*, *L. forbesi*, and *Ilex coindetii* show facultative semelparity, and, in the case of *S. officinalis*, a strikingly variable duration of reproduction (Boletzky, 1987, 1988; Baeg et al., 1993; Gonzalez, 1994; Gonzalez and Guerra, 1996; Rocha and Guerra, 1996; Melo and Sauer, 1999). Furthermore, in many of these species key traits—such as the timing and duration of reproduction—show substantial dependence on environmental effects (Rocha et al., 2001). Similar lability in these traits is also present in other clades, including both angiosperms and animals (see: Maltby and Calow, 1986; Tallamy and Brown, 1999; Hautekèete et al., 2001; Crespi and Teo, 2002; Varela-Lasheras and Van Dooren, 2014). Thus, because evolutionary change from one mode of parity to another is a ubiquitous life-history transition, accurately characterizing parity as a life-history variable of interest may prove crucial to accounting for the adaptiveness of life-history strategies.

## IV. UNDERSTANDING THE EVOLUTION OF PARITY AS A CONTINUOUS TRAIT

What changes should be made in light of the evidence that parity is a continuous trait? In this section I will focus on three main recommendations. First, I provide a short discussion of how seasonality should not be conflated with mode of parity. Second, I discuss the necessity of developing new mathematical modelling approaches that treat parity as a continuous variable. This is not simple, since parity itself is a composite trait, and relies on the coordination of many biological functions at once. Third, I discuss why ecologists should ground future studies of adaptive life history strategies in mechanistic details derived from genetic studies of continuously-varying life history traits underlying reproduction and, consequently, parity. These recommendations should improve both the validity and reliability of predictive models of life-history evolution, while simultaneously providing a framework for interpreting empirical findings regarding the expression of reproductive effort through time.

### (1) Seasonality and Mode of Parity

One major implication of treating parity as a continuous variable is that this reconception allows us to distinguish between parity and seasonality. Parity describes the concentration or diffusion of reproductive effort in time, which is distinct from the question of seasonal reproduction—i.e. how organisms should distribute reproductive effort among seasons, when seasonal cycles determine the favourability of establishment, growth, and reproductive conditions (Cole, 1954; Charnov and Schaffer, 1973; Schaffer, 1974b; Schaffer and Gadgil, 1975; Calow, 1979; Young, 1981; Bulmer, 1994; Evans et al., 2005; Ranta et al., 2007). It is, of course, clear that seasonality is related to parity. Insofar as an annual-semelparous organism is defined by the fact that it has a single reproductive episode that occurs within one year, it is likely to experience selection for strategies that optimize its reproductive schedule relative to season-specific environmental effects; this means that an annual-semelparous organism is more likely to show predictable seasonal patterns than a perennial-iteroparous congener that can escape a poor season by overwintering. However, the explanatory power of such seasonal adaptations may be much weaker when we compare a fast-reproducing semelparous organism with a slower-reproducing semelparous congener, or when we compare an iteroparous strategy where reproductive effort is distributed over two seasons with another where reproductive effort is distributed among ten seasons. Seasonal effects are likely to supervene on reproduction whenever regular intervals occur that have an impact on the favourability of reproduction. Thus it may be more fruitful to understand annuality and perenniality as strategies defined by the “digitization” of reproduction in response to seasonality. The advantage of this approach is that it makes it easier to understand flexible life histories, regardless of whether a species is semelparous or iteroparous.

There is widespread empirical evidence that seasonality and parity can vary independently. One common pattern is integer changes in voltinism among organisms that share a common mode of parity. For example, the Muga silkworm (*Antheraea assamensis*) is semelparous and multivoltine throughout its natural range (from India to Borneo). This species produces up to 6 generations per year, with the number of reproductive cycles depending on length of the season (Singh and Singh, 1998; Ghorai et al., 2009). However, the closely related Chinese tussar silkmoth, *Antheraea pernyi*, is bivoltine at the southern margins of its range, but is univoltine in northern China and Korea. Moreover, this continuous variation in voltinism along an ecological cline is due to continuous variation in environment-dependent biogenic monoamine production in the brains of diapause pupae (Fukuda, 1953; Matsumoto and Takeda, 2002; Liu et al., 2010). Life histories also vary continuously among populations of the wild silkmoth (*Bombyx mandarina*) and its domesticated counterpart (*Bombyx mori*), where populations in colder climates (e.g. European Russia) are univoltine, whereas those in China and Korea are bivoltine or multivoltine (Xia et al., 2009). Similar examples can also be found in crucifers (Williams and Hill, 1986; Springthorpe and Penfield, 2015), orchids (Chase et al., 2005), freshwater molluscs (Mackie and Flippance, 1983; McMahon and Bogan, 2001), and Centaurea (Asteraceae; Acker et al., 2014), among others. In each of these systems, a distinct continuum of reproductive strategies despite the supervening effect of seasonality is readily observable. Additionally, new models are being developed that consider generation length independently from parity (Waples, 2016). Thus we can easily tease apart the question of whether reproduction is concentrated in time—i.e. whether a given species is semelparous—from the question of whether seasonality requires that, in temperate climates, late-reproducing individuals should enter diapause rather than reproduce immediately.

### (2) Mathematical Models of Parity

A second problem facing life history theory is the challenge of making new mathematical models that account for the continuity of modes of parity. Although empirical studies of many taxa support the continuous conception of parity, the evolution of different modes of parity from one another has generally been explained by demographic models that compare the special case where annual-semelparous and perennial-iteroparous strategies have different demographic implications. This means that mathematical models that incorporate the assumption that there is a discrete difference between semelparous and iteroparous strategies may not be able to account for the real eco-evolutionary dynamics of the reproductive behaviour of many organisms that do not show either annual-semelparous or perennial-iteroparous modes of parity, or the reproductive behaviours of species or populations that display continuous differences in parity. To develop a general model that can account for all modes of parity as adaptive responses to environmental conditions should, as an axiom, treat parity as a continuous trait, and should be able to explain both the evolution of semelparous strategies from iteroparous ones (or vice versa) as well as the adaptive value of intermediate modes of parity. In this section I argue that dynamic mathematical models that integrate molecular data with demographic factors are the best candidates to explain the evolution of different modes of parity. In addition, extant mathematical models used to explain the adaptive differences between semelparous and iteroparous life histories—or to calculate “optimal” trait values for distinct semelparous-annual and iteroparous-perennial strategies—can and should be redesigned to incorporate a continuum of possible modes of parity.

These new models will have to build on and learn from a considerable body of existing models detailing the eco-evolutionary dynamics of semelparous and iteroparous life history strategies. Early conceptual and mathematical models of optimal semelparous reproduction were generally simple, deterministic, and were designed to predict a single “threshold” value that optimized life history characters such as size at first reproduction (Bell, 1980; Young, 1981). Threshold models of this type include senescence-threshold models based on the Penna aging model (Piñol and Banzon, 2011), as well as development-threshold models such as age-structured life history models. Age-structured models treat age at reproduction, and hence parity, as a discrete variable, and assess the evolutionary consequences of the degree of overlap between juvenile (i.e. prereproductive) and adult (reproductive) classes in a population (Wikan, 2012). Among the best known of these are Leslie models, which predict either few evolutionary stable states for semelparous organisms (Cushing, 2009, 2015; Cushing and Henson, 2012; Cushing and Stump, 2013) or even that populations should consist entirely of individuals of a given age class (Rudnicki and Wieczorek, 2014). Still other threshold models make similar predictions for survival traits (Da-Silva et al., 2008).

However, despite rare exceptions (e.g. Lesica and Young, 2005), threshold models generally fail to adequately predict reproductive phenotypes in field systems, prompting some authors to emphasize the importance of making fewer assumptions that contradict “real” life-history parameters—e.g. Burd et al. (2006) note that “empirical attention to norms of reaction across growth environments will be a more profitable approach than investigation of size thresholds per se”. That is, for short-lived semelparous species (and some long-lived semelparous species: see Rose et al., 2002), the impact of stochastic variation is substantial, and of equal or greater importance than the global optimum predicted by deterministic models (Rees et al., 1999). Furthermore, Rees et al. (1999) show that that deterministic age-structured models, which rest on the assumption that parity is discrete, consistently overestimate time at first reproduction in monocarpic plants. Empirical evidence showing a wide and highly variable range of reproductive phenotypes in natural populations (e.g. Marshall and Keough, 2007) has prompted the formulation of new modelling approaches that consider a range of semelparous strategies in response to environmental heterogeneity as a source of stochasticity that confounds threshold models that consider semelparous reproduction to be optimized for a given environmental (reviewed in Metcalf et al., 2003).

Recent mathematical models also fall into several types, each with a particular ecological focus. Integral projection models, which incorporate random fluctuations in environmental parameters related to reproduction, were developed to more accurately predict time to first reproduction and size at reproduction, both in iteroparous species (e.g. Kuss et al., 2008) and in semelparous species with a prolonged semelparous reproductive episode (e.g. Rees et al., 1999; Ellner and Rees, 2006 Rees et al. 1999; Sletvold 2005). Time-lagged models integral projection models attempt to account for the temporal discounting of reproductive value as well as size-specific effects on reproductive effort (Kuss et al., 2008). Newer age-structured stochastic models incorporate continuous variation in life history traits to predict optimal timing of reproduction; while these resemble earlier models that treat parity wholly as a discrete variable, the life-history traits in these models are treated continuously (Oizumi and Takada, 2013; Oizumi, 2014; Davison and Satterthwaite, 2016).

Several recent models have been developed to predict reproductive trait values given other (measured or measurable) life history parameters. This modelling methodology is intuitive and compatible with the idea that parity is a continuous trait. For example, Kindsvater et al (2016) used a stage-structured model to assess the degree to which trait covariation constrained life history adaptation in salmonids. Other kinds of data-driven models fall into two main types: (1) models that highlight the importance of phenotypically plastic reaction norms as maximizing fitness despite stochastic variability in environment (e.g. Burd et al., 2006); and (2) models that emphasize the innate variability in reproductive characters within species (e.g. Drouineau et al., 2014; Austen et al., 2015). Both of these ideas may be useful in modelling selective pressures on a continuum of modes of parity. Moreover, rather than using a single model to characterize semelparous investment in flowers and offspring, authors are now proposing a ‘meta-modelling’ approach to annual plant reproduction, recognizing that semelparous reproduction can be fine-tuned by natural selection through phenotypic plasticity (Hughes and Simons, 2014a). However, despite the success of some of these new models, as yet they still suffer from a relative paucity of data relative to other areas of life-history theory. When discussing models designed to predict the optimal timing of the initiation of reproduction, Metcalf et al. (2003) wrote that, “a glaring inconsistency between the [life history] models and the data is that all models predict a specific threshold flowering condition…but data from natural population show a graded response”.

One class of model that may prove to be useful in modelling the allocation of reproductive effort over time are dynamic state variable models (DSVMs). DSVMs are powerful dynamic optimization models used to characterize mechanistic relationships in ecology, and have the benefit of being able to be solved computationally (Clark and Mangel, 2000). Developing a DSVM can offer insight into the relative impact of underlying causal processes (in this case, the underlying patterns of genetic regulation of reproductive traits) on a state variable of interest (in this case, total plant fitness). Because the model follows the value of a state variable, the effects of multiple fitness components can be considered at once. Moreover, by parameterizing a DVSM with phenotypic data, ecologists can determine the additive and multiplicative contributions of variation at different gene loci, or between related phenotypes of interest. This is an important advantage insofar as continuous models of parity should, where possible include mechanistic detail. DSVMs are compatible with this approach: they can integrate a wide range of functional, spatial, structural, behavioural, or environmental limitations constraining investment in reproduction, and can generate testable predictions by determining optimal reproductive decision schedules (e.g. Yerkes and Koops, 1999; Skubic et al., 2004; Peterson and Roitberg, 2010).

### (3) Molecular Regulation of Parity

The third implication of understanding parity as a continuously varying trait is that further study of parity should be rooted in mechanistic detail, and identifying the mechanistic basis of different modes of parity (e.g. the contributions of individual genes and/or molecular pathways responsible for initiating and continuing reproduction) should be an important priority for evolutionary ecology. Integrating theoretical ecology with molecular biological data was not possible when early life-history models were developed, but since parity is determined by the onset and completion of reproductive episodes, and recent advances in molecular ecology have made it possible to understand the physiological and genetic basis of these events in many systems, this is, in many systems, an achievable goal. Numerous examples of continuously-expressed physiological processes result in continuous patterns of reproduction, and hence support the continuous conception of parity (Table 3). In this section, I will explain how parsing out the contributions of a single gene can improve our understanding of how modes of parity can vary continuously. To do so, I will discuss an important example: the control of the initiation of flowering in response to vernalization as it is regulated by FLOWERING LOCUS C (*FLC*) and its orthologues in the Brassicaceae. I discuss two such cases: (1) FLC regulation of vernalization response in *Arabidopsis thaliana*, a semelparous annual; and (2) PEP1 regulation of vernalization response in *Arabis alpina*, an iteroparous perennial.

**Table 3.**
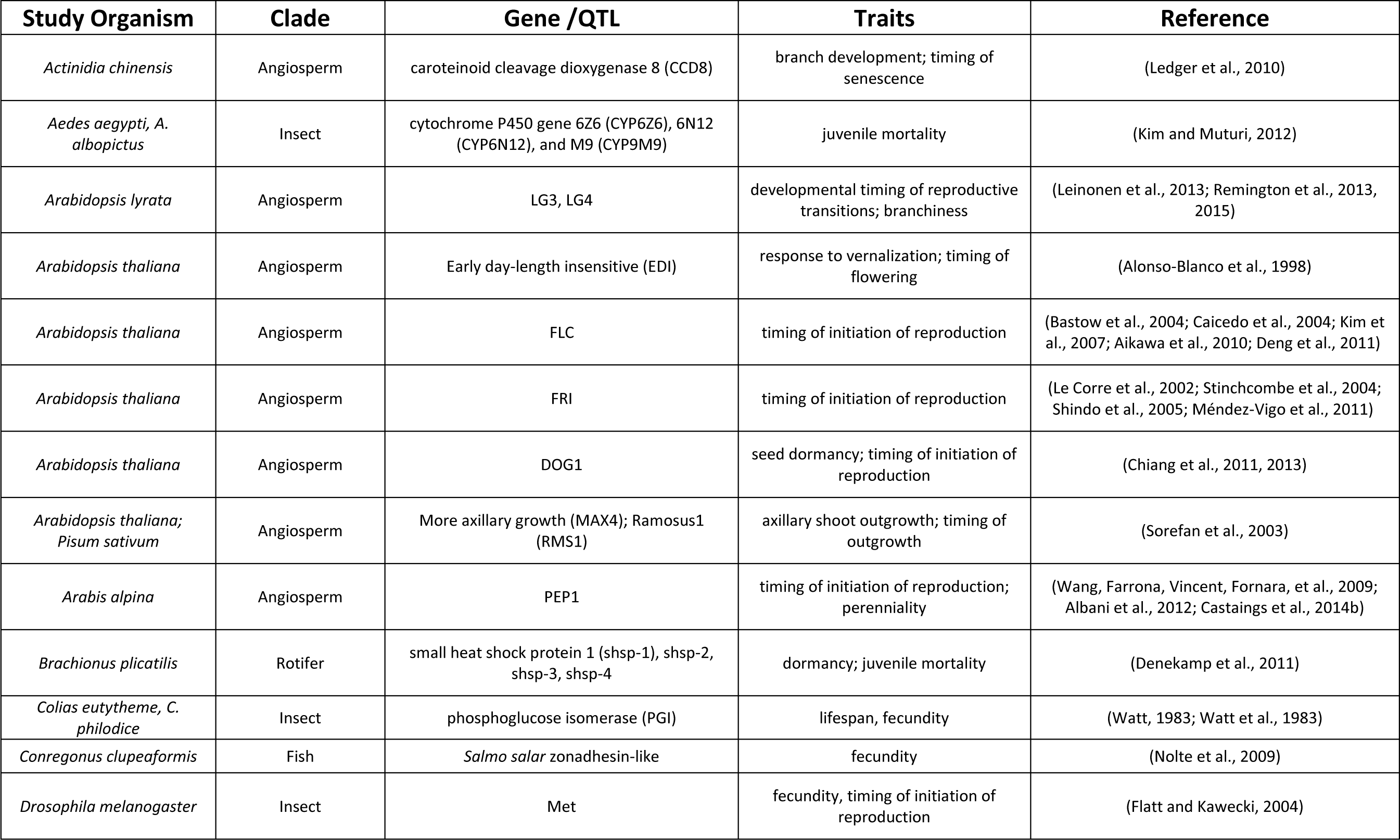
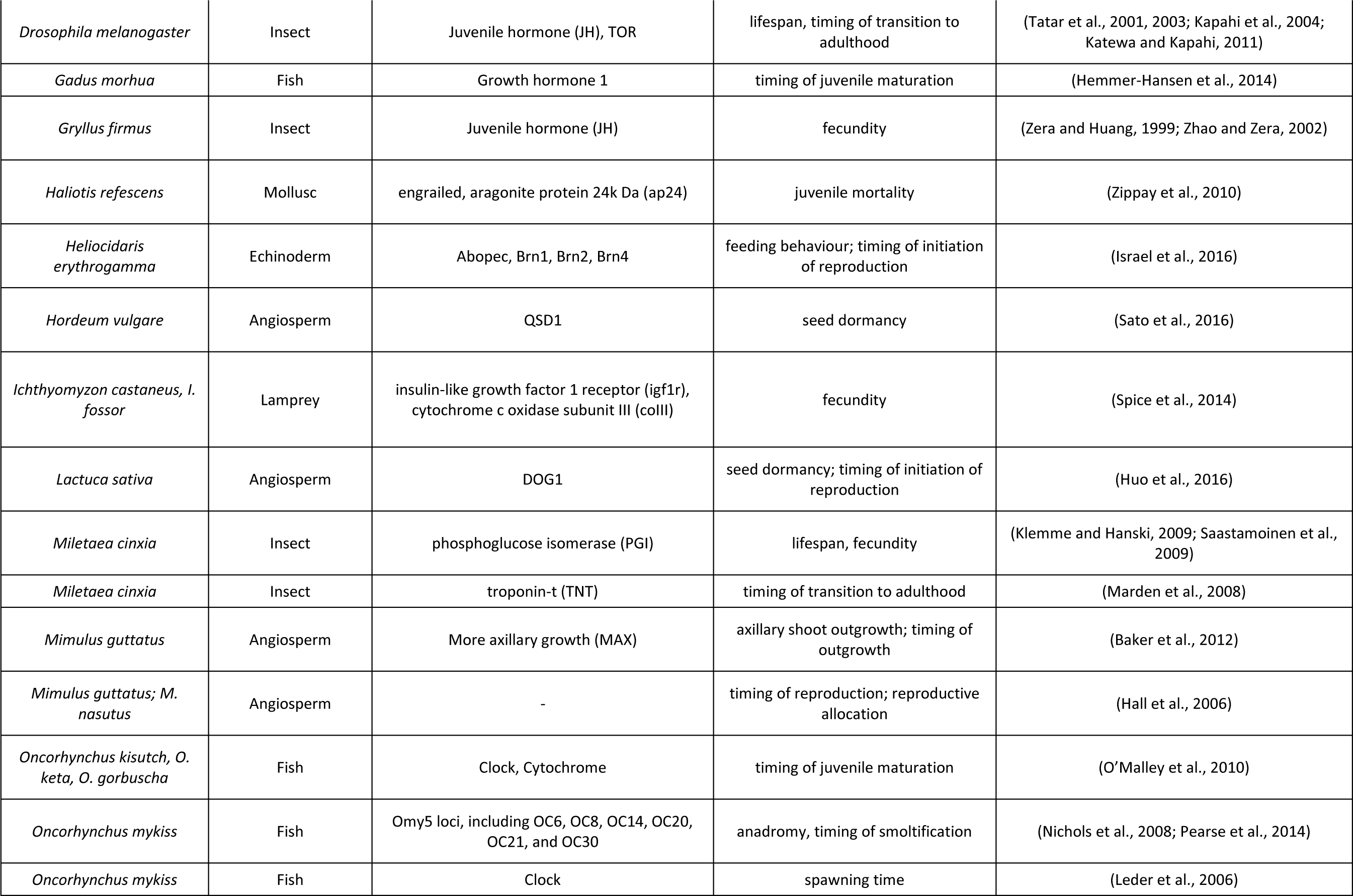
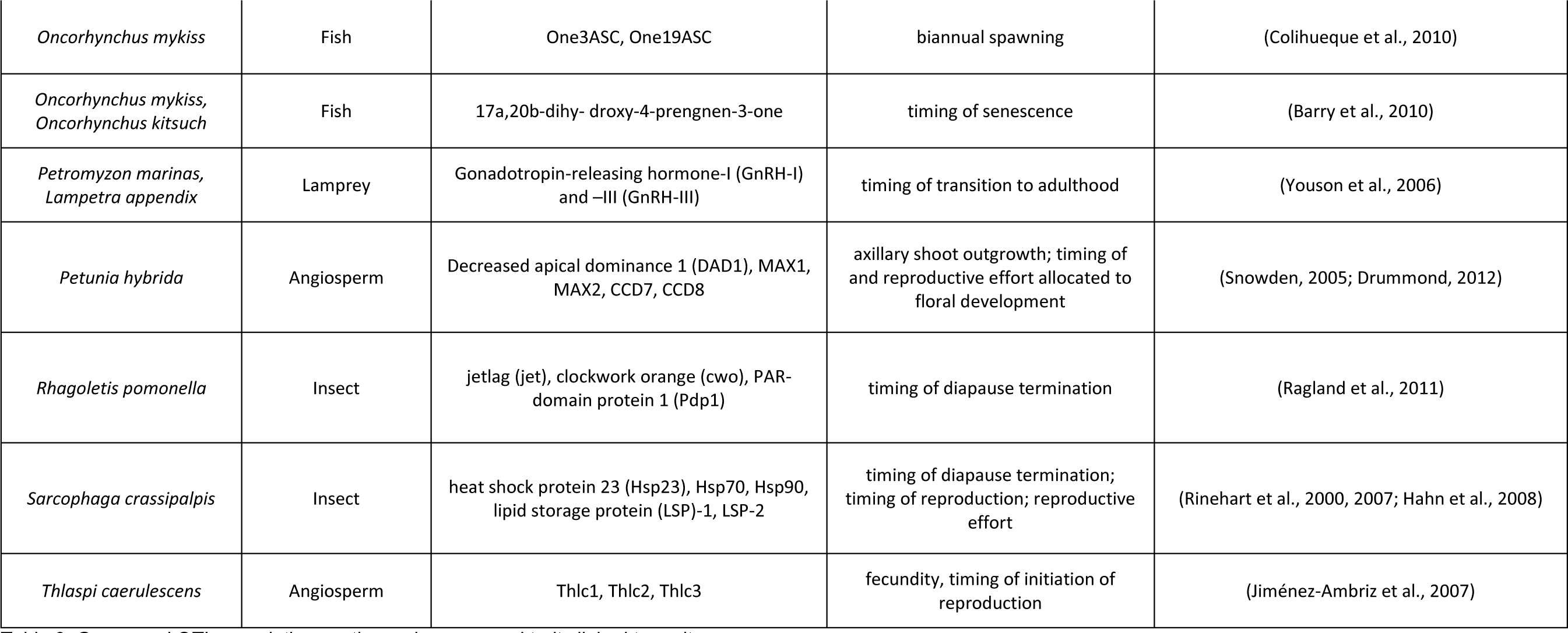
Genes and QTLs regulating continuously-expressed traits linked to parity.

| Study Organism | Clade | Gene /QTL | Traits | Reference |
| --- | --- | --- | --- | --- |
| *Actinidia chinensis* | Angiosperm | carotenoid cleavage dioxygenase 8 (CCD8) | branch development; timing of senescence | (Ledger et al., 2010) |
| *Aedes aegypti*, *A. albopictus* | Insect | cytochrome P450 gene 6Z6 (CYP6Z6), 6N12 (CYP6N12), and M9 (CYP9M9) | juvenile mortality | (Kim and Muturi, 2012) |
| *Arabidopsis lyrata* | Angiosperm | LG3, LG4 | developmental timing of reproductive transitions; branchiness | (Leinonen et al., 2013; Remington et al., 2013, 2015) |
| *Arabidopsis thaliana* | Angiosperm | Early day-length insensitive (EDI) | response to vernalization; timing of flowering | (Alonso-Blanco et al., 1998) |
| *Arabidopsis thaliana* | Angiosperm | FLC | timing of initiation of reproduction | (Bastow et al., 2004; Caicedo et al., 2004; Kim et al., 2007; Aikawa et al., 2010; Deng et al., 2011) |
| *Arabidopsis thaliana* | Angiosperm | FRI | timing of initiation of reproduction | (Le Corre et al., 2002; Stinchcombe et al., 2004; Shindo et al., 2005; Méndez-Vigo et al., 2011) |
| *Arabidopsis thaliana* | Angiosperm | DOG1 | seed dormancy; timing of initiation of reproduction | (Chiang et al., 2011, 2013) |
| *Arabidopsis thaliana*; *Pisum sativum* | Angiosperm | More axillary growth (MAX4); Ramosus1 (RMS1) | axillary shoot outgrowth; timing of outgrowth | (Sorefan et al., 2003) |
| *Arabis alpina* | Angiosperm | PEP1 | timing of initiation of reproduction; perenniality | (Wang, Farrona, Vincent, Fornara, et al., 2009; Albani et al., 2012; Castaings et al., 2014b) |
| *Brachionus plicatilis* | Rotifer | small heat shock protein 1 (shsp-1), shsp-2, shsp-3, shsp-4 | dormancy; juvenile mortality | (Denekamp et al., 2011) |
| *Colias eutytheme*, *C. philodice* | Insect | phosphoglucose isomerase (PGI) | lifespan, fecundity | (Watt, 1983; Watt et al., 1983) |
| *Conregonus clupeaformis* | Fish | *Salmo salar* zonadhesin-like | fecundity | (Nolte et al., 2009) |
| *Drosophila melanogaster* | Insect | Met | fecundity, timing of initiation of reproduction | (Flatt and Kawecki, 2004) |
| *Drosophila melanogaster* | Insect | Juvenile hormone (JH), TOR | lifespan, timing of transition to adulthood | (Tatar et al., 2001, 2003; Kapahi et al., 2004; Katewa and Kapahi, 2011) |
| *Gadus morhua* | Fish | Growth hormone 1 | timing of juvenile maturation | (Hemmer-Hansen et al., 2014) |
| *Gryllus firmus* | Insect | Juvenile hormone (JH) | fecundity | (Zera and Huang, 1999; Zhao and Zera, 2002) |
| *Haliotis refescens* | Mollusc | engrailed, aragonite protein 24k Da (ap24) | juvenile mortality | (Zippay et al., 2010) |
| *Heliocidaris erythrogamma* | Echinoderm | Abopec, Brn1, Brn2, Brn4 | feeding behaviour; timing of initiation of reproduction | (Israel et al., 2016) |
| *Hordeum vulgare* | Angiosperm | QSD1 | seed dormancy | (Sato et al., 2016) |
| *Ichthyomyzon castaneus, I. fossor* | Lamprey | insulin-like growth factor 1 receptor (igf1r), cytochrome c oxidase subunit III (coIII) | fecundity | (Spice et al., 2014) |
| *Lactuca sativa* | Angiosperm | DOG1 | seed dormancy; timing of initiation of reproduction | (Huo et al., 2016) |
| *Miletaea cinxia* | Insect | phosphoglucose isomerase (PGI) | lifespan, fecundity | (Klemme and Hanski, 2009; Saastamoinen et al., 2009) |
| *Miletaea cinxia* | Insect | troponin-t (TNT) | timing of transition to adulthood | (Marden et al., 2008) |
| *Mimulus guttatus* | Angiosperm | More axillary growth (MAX) | axillary shoot outgrowth; timing of outgrowth | (Baker et al., 2012) |
| *Mimulus guttatus; M. nasutus* | Angiosperm | - | timing of reproduction; reproductive allocation | (Hall et al., 2006) |
| *Oncorhynchus kisutch, O. keta, O. gorbuscha* | Fish | Clock, Cytochrome | timing of juvenile maturation | (O'Malley et al., 2010) |
| *Oncorhynchus mykiss* | Fish | Omy5 loci, including OC6, OC8, OC14, OC20, OC21, and OC30 | anadromy, timing of smoltification | (Nichols et al., 2008; Pearse et al., 2014) |
| *Oncorhynchus mykiss* | Fish | Clock | spawning time | (Leder et al., 2006) |
| *Oncorhynchus mykiss* | Fish | One3ASC, One19ASC | biannual spawning | (Colihueque et al., 2010) |
| *Oncorhynchus mykiss*,<br>*Oncorhynchus kitsuch* | Fish | 17a,20b-dihydroxy-4-pregnene-3-one | timing of senescence | (Barry et al., 2010) |
| *Petromyzon marinus*,<br>*Lampetra appendix* | Lamprey | Gonadotropin-releasing hormone-I (GnRH-I)<br>and -III (GnRH-III) | timing of transition to adulthood | (Youson et al., 2006) |
| *Petunia hybrida* | Angiosperm | Decreased apical dominance 1 (DAD1), MAX1,<br>MAX2, CCD7, CCD8 | axillary shoot outgrowth; timing of<br>and reproductive effort allocated to<br>floral development | (Snowden, 2005; Drummond, 2012) |
| *Rhagoletis pomonella* | Insect | jetlag (jet), clockwork orange (cwo), PAR-domain protein 1 (Pdp1) | timing of diapause termination | (Ragland et al., 2011) |
| *Sarcophaga crassipalpis* | Insect | heat shock protein 23 (Hsp23), Hsp70, Hsp90,<br>lipid storage protein (LSP)-1, LSP-2 | timing of diapause termination;<br>timing of reproduction; reproductive<br>effort | (Rinehart et al., 2000, 2007; Hahn et al., 2008) |
| *Thlaspi caerulescens* | Angiosperm | Thlc1, Thlc2, Thlc3 | fecundity, timing of initiation of<br>reproduction | (Jiménez-Ambríz et al., 2007) |

In *Arabidopsis*, continuous variation in parity–i.e. the timing of floral initiation and the duration of flowering—is determined by continuous expression of flowering-time genes, including *FLOWERING LOCUS T* (*FT*) (Kardailsky, 1999; Yanovsky and Kay, 2002; Kotake et al., 2003; Imaizumi and Kay, 2006; Turck et al., 2008; Simon et al., 2015), *FRIGIDA* (*FRI*) (Johanson et al., 2000; Le Corre et al., 2002; Michaels et al., 2004; Stinchcombe et al., 2004; Shindo et al., 2005; Schläppi, 2006), *FLOWERING LOCUS C* (Coupland, 1995; Amasino, 1996; Michaels and Amasino, 1999; Sheldon et al., 2000; Michaels et al., 2003, 2004; Bastow et al., 2004; Imaizumi and Kay, 2006; Kim et al., 2007; Chiang et al., 2009), *GIGANTEA* (*GI*) (Fowler et al., 1999; Mizoguchi et al., 2005; Jung et al., 2007), and *CONSTANS* (*CO*) (Redei, 1962; Putterill et al., 1995; Koornneef et al., 1998; Samach et al., 2000; Suárez-López et al., 2001; Valverde et al., 2004). Through different pathways, *GI* and *CO* activate the floral integrator gene *FLOWERING LOCUS T* (*FT*), which transcribes a protein that activates floral identity genes in the shoot apical meristem (Turck et al., 2008; Tiwari et al., 2010). In contrast, *FLC*—along with *FRI*, which regulates *FLC* transcription—represses flowering in the absence of vernalization.

Although much is known about the complex question of how flowering is induced, here I concentrate on *FLC*, since continuous differences in *FLC* expression cause continuous variation in the duration and timing of semelparous reproduction (Wilzcek et al., 2009; Burghardt et al., 2015). This variation causes continuous differences in parity. For instance, throughout Europe, parity in wild populations of *Arabidopsis* seems strongly determined by climate and/or latitude: toward the colder margins of its range, in northern Finland, plants show a fast-cycling summer semelparous annual life history, while populations near the Mediterranean show a winter annual life history, and populations in intermediate locations (e.g. the UK) display intermediate life histories (Thompson, 1994; Méndez-Vigo et al., 2011; Ågren and Schemske, 2012). Lab studies have identified *FLC* as the mechanism responsible for this life-history variation. For example, Wilzcek et al. (2009) introgressed a functional *FRI* allele into *A. thaliana* ecotypes with nonfunctional alleles. They predicted that this genetic modification— which causes the upregulation of *FLC*—would see plants transition from a summer-annual to winter-annual life history. Instead, plants with functional *FRI* alleles flowered only 10 days later than those with nonfunctional *FRI* alleles, causing the authors to note that their results “suggest that *A. thaliana* ecotypes cannot simply be divided into two discrete classes of winter-annual and rapid-cycling genotypes. Rather, most ecotypes may be capable of both life histories” (p. 933). This prediction is consistent with recent data from studies of the impact of *FLC* on the life histories of *A. thaliana* ecotypes sourced from different parts of its native range. While populations varying at the *FLC* locus show substantial local adaptation with respect to important life history traits—including those, such as length of duration of reproduction, which underlie mode of parity—most ecotypes adopt new life histories when translocated to radically different environments (Ågren and Schemske, 2012; Dittmar et al., 2014; Ågren et al., 2016; Postma and Ågren, 2016).

Where the prevalence of different *FLC* alleles differs between populations, differential expression of FLC can result in different flowering phenologies, and even different modes of parity (Johanson et al., 2000; Michaels et al., 2004; Schläppi, 2006; Banta and Purugganan, 2011). This is likely an adaptive response; life-history models of the natural genetic variation at the *FLC* and *FRI* loci have shown that *FLC* expression explains a relatively high level of variation in fitness (Donohue et al., 2014; Burghardt et al., 2015; Springthorpe and Penfield, 2015). Moreover, empirical studies suggest that such fitness differences may account for the latitudinal cline in *Arabidopsis* life history found in natural populations (Caicedo et al., 2004). Thus it seems that local adaptation of different modes of parity result from populations experiencing stabilizing selection for climate-appropriate *FLC* alleles (Postma and Ågren, 2016).

While fitness differences are tightly linked to phenotypic variation, major plant phenotypes such as flowering phenology are highly plastic in *Arabidopsis*. The genetic and epigenetic regulation of *FLC* regulate the timing of important life-history transitions, determining a plant’s reproductive schedule, and hence its mode of parity (Albani and Coupland, 2010; Turck and Coupland, 2014). However, environmental factors such as seed maturation and germination temperatures can interact with plant genotypes to produce “clusters” of phenotypically similar, yet genotypically distinct, plant phenologies (Burghardt et al., 2016). Environmental variation can therefore facilitate the adoption of multiple flowering phenologies (e.g. summer annual, winter annual, rapid cycling, etc.)— and thus modes of parity—from a single *Arabidopsis* ecotype (Simpson and Dean, 2002; Méndez-Vigo et al., 2011).

The relationship between *FLC* and variation in mode of parity is best understood in *Arabidopsis*, continuous variation in parity arising from response to vernalization has also been identified in the confamilial species *Arabis alpina*. *PERPETUAL FLOWERING 1* (*PEP1*), the orthologue of *FLC* in *A. alpina* (Wang, Farrona, Vincent, Joecker, et al., 2009), is responsible for regulating vernalization response. In *Arabidopsis*, *FLC* confers a obligate vernalization requirement on individuals; thus flowering is not possible until plants experience winter temperatures, which causes chromatin remodeling that prevents *FLC* transcription (Sheldon et al., 2000; Bastow et al., 2004). *PEP1* also confers an obligate vernalization requirement to *A. alpina*, however this effect is (seasonally) temperature-dependent. That is, in the Pajares ecotype, *PEP1* transcription is temporarily repressed by low temperatures, mediated by the continuous expression of *COOLAIR*-family antisense RNAs produced in response to cold (Wang, Farrona, Vincent, Joecker, et al., 2009; Albani et al., 2012; Castaings et al., 2014a), and thus facilitating the regular alternation of vegetative and reproductive phases that characterize iteroparous-perennial life histories. Like *FLC*, considerable natural variation exists at the *PEP1* locus—most natural *A. alpina* ecotypes have a functional *PEP1* allele, and under controlled conditions restrict flowering duration, although some accessions lack a functional *PEP1* allele, and thus continuously flower without vernalization (Torang et al., 2015). That *PEP1* alone is responsible for this phenotype has been confirmed using mutant lines (Wang, Farrona, Vincent, Joecker, et al., 2009; Wang et al., 2011; Albani et al., 2012; Bergonzi et al., 2013). However, this transition is continuous rather than discrete, since the degree of *PEP1* suppression depends both on *PEP1* allele present in the accession as well as the duration of vernalization.

Thus, in both *Arabidopsis thaliana* and *Arabis alpina* (among other Brassicaceae spp.), parity is linked to genetic variation at the FLC/PEP1 locus, and alleles from different, locally-adapted ecotypes show a range of distinct phenotypes (Aikawa et al., 2010; Kemi et al., 2013; Zhou et al., 2013). Moreover, FLC itself is subject to a number of regulatory mechanisms, some of which result in independent life history differences (Gazzani et al., 2003; Shindo et al., 2005, 2006). *FLC* example is not an isolated case, nor have genes linked closely to parity been discovered only in plant species; although a comprehensive description of all genes linked to parity in all species is beyond the scope of this article, a few notable examples from a variety of well-studied taxa are presented in Table 3. Genes linked to traits underlying parity, including reproductive maturation, stress response, reproductive phenology, and senescence, have been the subject of many informative reviews (see: McBlain et al., 1987; Finch and Rose, 1995; Tower, 1996; Danon et al., 2000; Eulgem et al., 2000; Rion and Kawecki, 2007; Garcia De Leaniz et al., 2007; Hall et al., 2007; Schneider and Wolf, 2008; Xin et al., 2008; Costantini et al., 2008; Amasino, 2009; Thomas et al., 2009; Partridge, 2010; McCormick et al., 2011; Selman and Withers, 2011; Kenyon, 2011; Thomas, 2013; Blümel et al., 2015; Wang et al., 2015).

## V. CONCLUSIONS

We still know far too little about why the evolutionary transition from semelparity to iteroparity (or vice versa) is as common as it is, or under which ecological conditions intermediate strategies—such as facultative semelparity—will thrive. While models rooted in the conception of parity as a binary trait do a good job of accounting for the fitness differences between discrete semelparous-annual and iteroparous perennial alternative strategies, systems characterized by only these possibilities—and no others— are special cases. In many cases the life history question at hand is more subtle: why does a given species evolve a facultative strategy, or why does another show intraspecific variation in the length of the semelparous reproductive episode? In such cases—which are not as rare as they were when the adaptiveness of parity was first being investigated— the discrete conception of parity is a impractical oversimplification.

Thus the main conclusion of this work is that parity should be treated as a continuous trait rather than a discrete one. This reconception of parity offers several notable advantages for life-history theory. First, treating parity as a continuous trait allows us to treat parity as a distinct life history syndrome, itself the result of correlated selection on a suite of traits, that may show finely-graded correlated variation within species or populations. This is advantageous because parity is a composite trait, and the act of reproducing at a given time, of a given duration, etc. involves the recruitment and coordination of many independent parts, each of which may affect the expression of others. An integrative approach has proven to be valuable in studying other multifactorial composite traits (e.g. Buoro and Carlson, 2014). Furthermore, whether they share common genetic basis or not, obvious or visible life history characters may not be primary targets of selection, and evolution of such traits may occur as an epiphenomenon of selection on (one or many) other apparent or non-apparent underlying traits. Second, developing accurate mathematical and mechanistic models designed to explain fitness differences between life histories—as well as to understand the nature of the costs and trade-offs associated with initiating and completing reproductive episodes—will be best understood by acknowledging that living systems show a continuum of reproductive strategies between annual-semelparity and perennial-iteroparity. Considering only annual-semelparity and perennial-iteroparity as discrete alternatives, although a useful simplification for many models, is biologically accurate only in a limited number of special cases, and the continuous conception of parity is more likely to approximate the eco-evolutionary dynamics of natural systems that show intraspecific or plastic variation in the expression of parity. Finally, treating parity as a continuous variable that represents a syndrome of associated traits may make it easier to integrate life history studies with mechanistic details deriving from molecular ecology, insofar as composite life history traits such as parity are unlikely to be the result of a simple presence or absence of a single gene or allele. Instead, parity is likely to be the product of complex systems of genetic, translational, and post-translational regulation.

There are important issues to consider for further work on a continuum of modes of parity. First, life-history theorists should make their language clearer and tighten definitions, in order to avoid confusion and improve the effectiveness of evolutionary explanations. Parity, the degree of temporal concentration or diffusion of reproductive effort in time, is a distinct evolutionary question from whether organisms reproduce in one year or in more than one year (“classic” semelparity and iteroparity), or whether or not organisms have a distinct life cycle that is completed according to an annual cycle or not (annuality and perenniality). Consider two annual plants as an example: the concentration of reproductive effort in time—i.e. realizing a hyper-semelparous (or uniparous) reproductive strategy—is a different strategy than spreading reproduction out over many months, even if both strategies are completed within a single year. Moreover, the evolutionary and mechanistic reasons that account for seasonal behaviours—e.g. annuality and perenniality—are distinct from those that account for the degree of parity. Making these distinctions clear, both in language and in predictive models, should be an important priority for evolutionary ecologists interested in the problem of parity.

Other important issues exist as well. A second question is that the extant body of theory—much of which describes various special cases in which demographic or environmental factors confer a fitness advantage on an annual-semelparous strategy over a perennial-iteroparous one (or vice versa)—should be preserved and reworked into a more general explanation of how factors affecting fitness through time determine optimal patterns of the distribution of risk in and among seasons. More general models should consider a range of possible modes of parity. A third issue is that reproductive characters of long-lived semelparous species are generally easier to model than characters of short-lived species, and while environmental heterogeneity plays an important role determining the optimal allocation of reproductive effort in annual semelparous species, long-lived semelparous species can afford to be “choosier” about when they reproduce, and therefore have been shown to more closely approximate model predictions. This may be especially true when, as in many long-lived perennial-iteroparous species, the relationship between age and cost of reproduction is nonlinear. Thus, developing models that accurately model the fitness dynamics of short-lived semelparous species should be a priority. Fourth, intraspecific differences in life-history strategy, particularly between populations inhabiting dissimilar environments, can confound the degree to which empirical studies can test predictions of *a priori* models or parameterize them. Additional empirical comparisons of different modes of parity should therefore be as general as possible, and incorporate as many points along the continuum of modes of parity as possible, in order to maximize generalizability. Last, new models of modes of parity should, on a system-by-system basis, attempt to account for the underlying mechanisms that determine the timing and nature of the allocation of reproductive effort. Because modes are composite traits, integrating the molecular mechanisms that underlie many traits into a single model is not a simple task. Moreover, even orthologous genes with may have dramatically different effects in closely related species, and this unpredictability may considerably complicate the process of mechanistically modelling of modes of parity. Despite the considerable challenge that it poses, integrating molecular ecological studies of the fitness consequences of different genotypes with theoretical mathematical models should be an important priority in the medium-to long-term future, and would represent an important area of consilience between molecular and theoretical ecology.

## VI. ACKNOWLEDGEMENTS

I would like to thank Andrew Simons, Maria Albani, Wim Soppe, and Amanda Feeney for their assistance in developing the ideas in this manuscript. This work was supported by a Postdoctoral Fellowship from the Natural Sciences and Engineering Research Council and a Humboldt Research Fellowship from the Alexander von Humboldt Foundation.

